# The impact of *P*-Element-induced hybrid dysgenesis on the male germline in *Drosophila simulans*

**DOI:** 10.64898/2026.06.28.735054

**Authors:** Joanne S. Griffin, Ewan Harney, Charlton Capes, Rowan Connell, Andrea J. Betancourt, Valeria Romero-Soriano

## Abstract

The P-element, a DNA transposon, has independently invaded two *Drosophila* species, accompanied by rapid evolution of suppression. In the germline, suppression is mediated primarily by maternally expressed piRNAs, a class of regulatory small RNAs associated with PIWI proteins. The offspring of females that lack P-element-specific piRNAs and males that contain P-elements suffer a syndrome of deleterious phenotypes, including sterility, genome rearrangements, gonadal atrophy, and mutations, while the offspring of the reciprocal cross are normal. These effects, collectively termed hybrid dysgenesis, have been investigated primarily in female *D. melanogaster*. Here, we study hybrid dysgenesis in male *D. simulans*. Using an attached-X chromosome stock, we generated genetically identical F1 males that differed only in maternal suppression of the P-element. Using targeted sequencing of P-element breakpoints, we show that P-element transposition is elevated in dysgenic males and confirm a preference for insertion near origins of replication. Using transcriptomics, we show that dysgenic males have elevated P-element expression and reduced splicing suppression, with patterns of gene expression suggesting the loss of mature sperm cells. Fertility assays show higher rates of male sterility but otherwise modest effects on fertility. In conjunction with the transcriptomic data, small RNA sequencing confirms that the piRNA pathway functions in testes. Our results suggest that the P-element may spread more readily through males than females, as transposition rates are similar while fertility defects are less severe in males.

**Summary:** Transposable elements (TEs) are selfish genes that replicate independently of host genomes. Hosts produce piRNA molecules to suppress established TEs, but for new TEs, piRNAs are not available. This has strong negative consequences for females, but effects in males are unclear. The fly *Drosophila simulans* has recently acquired a TE, the P-element, and males from lines lacking P-element piRNAs suffer more P-element expression and replication than controls, leading to reduced expression of genes associated with mature sperm cells, and sometime sterility. Yet unsterilised males did not suffer fertility loss, suggesting that TEs may spread more easily in males than females.

## Introduction

Transposable elements (TEs) are selfish DNA sequences found in nearly all eukaryotic species. They propagate by inserting into new locations in a genome, potentially causing deleterious or beneficial mutations (reviewed in Betancourt et al., 2024; Biémont, 2010; McDonald, 1993). Because they often replicate more quickly than they are lost, TE copies can accumulate in the genome over time, with both functional and degraded copies sometimes contributing substantially towards genome size (Gregory 2005; Canapa et al. 2015; Marino et al. 2025). TEs also spread *via* horizontal transfer, in which individuals acquire DNA from unrelated organisms (Schaack et al. 2010). If newly invading TEs succeed in inserting themselves into the germline, they have the potential to be passed on vertically to offspring and spread in a population. Whilst the mechanism of transfer is often unknown, horizontal transfer of TEs in insects appears to be common (Bartolome et al. 2009; Peccoud et al. 2017; Scarpa et al. 2024). One of the best-known examples of this is the P-element, which invaded *Drosophila melanogaster* from a distantly related species, *D. willistoni*, in the mid-20^th^ century (Daniels et al. 1990). The P-element also subsequently invaded the sibling species of *D. melanogaster*, *D. simulans*, spreading with remarkable speed (Kofler *et al*, 2015; Hill *et al*, 2016).

The original discovery of the P-element was prompted by the detection of abnormal traits in the offspring of crosses between freshly collected wild and old laboratory strains of *D. melanogaster*, eventually found to be due to a 2.9kb DNA transposon present only in wild strains (Hiraizumi 1971; Kidwell et al. 1977; Bingham et al. 1982). Specifically, when wild males that have the P-element are crossed to females from laboratory strains that lack the P-element, their offspring suffer from aberrant traits such as malformation of the gonads, male recombination, increased mutation rate and sterility– traits collectively termed ‘hybrid dysgenesis’ (Kidwell et al. 1977; Kidwell 1983). Though most studies of hybrid dysgenesis in *Drosophila* focus on females, the phenotype was first described in males, where it appears to be relatively mild (Hiraizumi 1971; Kidwell et al. 1977). Similarly, the piRNA pathway (Vagin *et al*, 2006) –the small RNA system that is the major suppressor of TEs in the germline (Aravin et al. 2001) – also shows relatively low expression in males (Saint-Leandre et al. 2020). This raises the possibility that spread of the P-element may occur more easily through F1 males than through females: new insertions would be more likely to passed on through males, which are less affected by sterility.

Here, we investigate several aspects of the dysgenesis phenotype in males, including fertility, piRNA expression, and abnormal gene and TE expression in the testes of *D. simulans* males. Following Engels (1979), we take advantage of an attached-X stock to obtain F1 males with identical nuclear genomes from reciprocal crosses, allowing us to compare males that lack maternal P-element suppression with genetically identical controls. We find that these males show elevated expression of the P-element in their testes, and, along with their sisters, higher rates of P-element transposition than reciprocal controls. Accordingly, these males also appear to suffer germ-cell death and mildly reduced fertility.

## Materials and methods

### Fly stocks and culturing

*D. simulans* lines used in this study were the ST8/C(1)RM,yw attached-X stock (Cazemajor et al. 2000), M252 (collected in Madagascar; Palmieri et al., 2015), and Cro18 (collected in Croatia in 2014 by A. Jakšić; Hill et al., 2016). M252 and the attached-X stock both lack the P-element, while Cro18 contains P-element insertions and can induce dysgenesis (Hill et al. 2016). Females of the attached-X stock carry a compound-X chromosome (consisting of two full-length X chromosomes sharing a centromere) and a Y-chromosome (X^X/Y), while males have a normal X/Y karyotype. A substrain of the attached-X line was used to ensure homogenisation of the Y-chromosome. Flies were maintained on a cornmeal-based diet (ASG) of yeast, sugar, maize, nipagin and propionic acid at 25°C under a 12 h:12 h light:dark cycle, unless stated otherwise.

### Fly crosses

P-type lines contain P-element and their maternal suppressors; M-type lines lack both. Crosses between P-type males and M-type females induce dysgenic phenotypes in the offspring, while the reciprocal cross results in normal offspring. Normal male offspring of reciprocal crosses differ in the parental origin of their sex chromosomes. To obtain dysgenic and non-dysgenic (reciprocal) F1 males with identical nuclear genotypes, we made use of the attached-X stock, reciprocally crossed to the P-type Cro18 strain as shown in Figure 1. Crosses were performed and offspring were reared at high temperature, when dysgenesis effects are strongest (28.5°C). For the transposition assay, we also obtained dysgenic and non-dysgenic F1 females by performing reciprocal crosses between M252 and Cro18 at 28.5°C.

**Figure 1.**
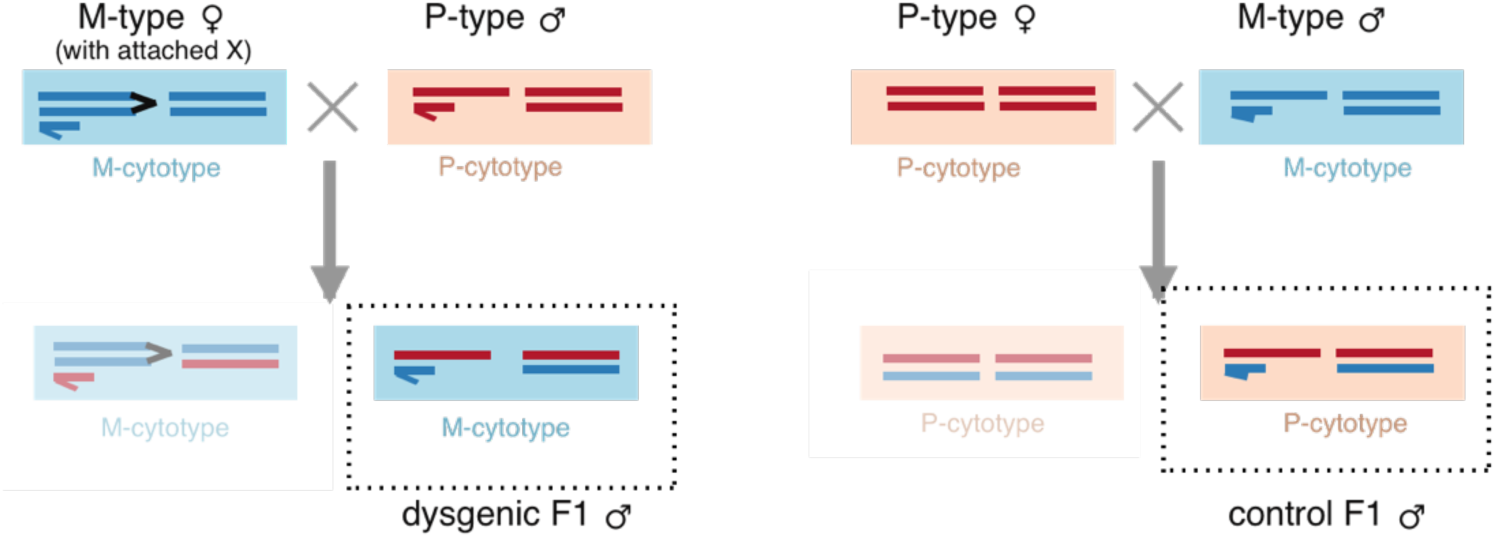
Reciprocal MxP crosses to obtain dysgenic and reciprocal control F1 males with identical nuclear genotypes and contrasting cytotypes. The karyotype is represented by bars with the sex-chromosomes on the left, and a representative pair of autosomes on the right; cytotype is represented by the coloured background of each genotype. The colour indicates origin in the M- or P-cytotype stock. The M-cytotype, attached-X females carry a pair of X-chromosomes that segregate together and a free Y chromosome; males of this stock have a normal XY genotype. In crosses between X^^^XY females and XY males, the usual pattern of sex-chromosome inheritance is reversed, with sons inheriting X-chromosomes from their fathers, and Y-chromosomes from their mothers (left) The reciprocal cross has normal sex-chromosome inheritance (right). Males used for the comparisons are shown in dashed boxes.

### Fertility assay

Dysgenic and non-dysgenic males were individually crossed to P-type Cro18 virgin females at 25°C, with two females provided per male. Males were crossed on the day they emerged. We performed a total of 56 crosses (26 with dysgenic and 30 with non-dysgenic F1 males). We provided fresh food vials every 2-3 days, replacing females if dead. Adults emerging from the vials were sexed and counted every 2-3 days for 39 days or until no more offspring emerged.

### RNA extraction, library prep and sequencing

We dissected testes from the parental stocks (attached-X and Cro18) at 2-4 days old; as well as from the F1 offspring from both dysgenic and reciprocal crosses, at two time points: 2-4-days old (herein referred to as young flies) and 21-days old (herein referred to as old flies). Dissected testes were frozen using liquid nitrogen and stored at −80°C prior to RNA extraction. Total RNA was extracted from pooled gonads of ∼10 flies using the NucleoSpin RNA purification kit (Macherey-Nagel, Düren, Germany) for RNA-seq and using TRIzol Reagent (Invitrogen, Waltham, MA, USA) for small RNAseq. RNA was extracted from two biological replicates for each sample combination of age and genotype (16 samples in all), then RNA concentration was measured with Qubit (Thermo Fisher Scientific, Waltham, MA, USA), and RNA integrity checked on a Bioanalyzer (Agilent, Santa Clara CA, USA).

RNA-seq libraries were prepared at the Centre for Genomic Research (CGR, Liverpool), using the riboPOOL rRNA depletion kit (*D. melanogaster*) followed by the NEBNext Ultra II Directional RNA library prep kit (New England Biolabs, Ipswich, MA, USA). RNA-seq libraries (two replicates per sample) were then sequenced on one lane of an SP flow cell on the Illumina NovaSeq 6000 (Illumina, San Diego, CA, USA). Raw reads were trimmed for the presence of Illumina adapter sequences using Cutadapt version 1.2.1, with parameter -O 3 (Martin, 2011) and then further trimmed using Sickle version 1.2 (Joshi and Fass, 2011) with a minimum window quality score of 20.

Prior to small RNA-seq library prep (CGR, Liverpool), a DNAse treatment was performed on the TRIzol-extracted samples. Libraries were then prepared using the NEBNext small RNA library prep kit, including an additional 2S rRNA blocking step (as described in Wickersheim & Blumenstiel, 2013). Those libraries were also sequenced on one lane of an SP flow cell on the Illumina NovaSeq 6000.

### Expression analysis– genes

We quantified transcript abundance using the mapping-based mode of Salmon v1.10.2 (Patro et al. 2017), quantifying reads for each sample against the *D. simulans* reference transcriptome (version GCF_016746395.2_Prin_Dsim_3.1, NCBI). We generated transcript count and expression matrices by importing data into R with tximport (Soneson et al. 2016). We then removed genes that had a count of 10 or less. A differential gene expression analysis of reciprocal and dysgenic, young and old flies was performed on the counts matrix using DESeq2 (Love et al. 2014) in R, with the FDR set to 0.05.

We conducted a gene ontology (GO) over-representation analysis using the R package TopGO (Alexa et al., 2006; Alexa and Rahnenführer, 2024), retrieving annotated GO terms from the AnnotationHub R package (Morgan and Shepherd, 2024) using the most specific GO terms to create a gene-to-GO map, and the readMappings function to parse the gene set. GO terms with a *P* < 0.01 from a Fisher’s exact test were considered significantly enriched. We investigated over- and under-representation of cell types in reciprocal and dysgenic testes by comparing our differentially expressed genes to annotations and cell-type markers from single-cell sequencing of *D. melanogaster* testes (Raz et al. 2023) and using EWHA (Skene and Grant 2016) to test for significant under- or over-representation of known cell types.

### Expression analysis– transposable elements and small RNAs

We quantified TE abundance from the RNA-seq reads using SalmonTE, run using the parameters --exprtype count –num, and the *D. simulans* Prin3.1 reference genome. We used the TE library from Chakraborty et al. (2021), filtered of satellites, rRNA and low complexity sequences, with the addition of the *D. simulans* P-element (NCBI accession KP256109.1), plus 12 additional transposable element sequences (Scarpa et al. 2024). We treated the small RNAs as above, after filtering for read lengths of 23-32 bp which correspond to putative piRNAs. We used only R1 reads (which are in the same sense/anti-sense orientation as the original molecules) from these paired end sequencing runs, as the lengths of the reads exceed those of the small RNA molecules. Again, we generated transcript count and expression matrices by importing data into R and removed genes that had a count of 10 or less. A differential gene expression analysis of reciprocal and dysgenic, young and old flies was performed on the counts matrix using DESeq2 (Love et al. 2014) in R, with the FDR set to 0.05. To test for the 10 bp overlap characteristic of ping-pong amplification in piRNAs, we used the signatures.py script of Antoniewski (2014), available at https://github.com/ARTbio/tools-artbio/blob/main/tools/small_rna_signatures/signature.py, run with parameters --minquery 23 --maxquery 34 --mintarget 23 --maxtarget 34 --minscope 1 --maxscope 19.

### Splicing regulation

As the P-element can be regulated by suppression of splicing, we tested for an elevated splicing rate in dysgenic flies compared to reciprocal controls. We estimated the number of spliced and unspliced P-element transcripts in both types of F1 males. We filtered for RNAseq reads mapping to the P-element, discarded singleton reads with repair.sh (Bushnell, 2016; https://sourceforge.net/projects/bbmap/) and remapped these reads to an annotated P-element reference using STAR (v2.7.11b). We tested specifically for splicing of IVS3 (the third P-element intron), as splicing suppression targets this intron particularly (Laski et al, 1986). We estimated the number of spliced transcripts using the SJout.tab output of STAR to count reads crossing splice junctions. We estimated unspliced transcripts from mapped reads that overlap with IVS3 intron and flanking exons using bedtools (v2.31.1, Quinlan and Hall, 2010) intersect, using 10nt on either side of each intron-exon junction. Inspection of the data revealed that some transcripts used alternate donor/acceptor sites, resulting in a splice variant including a small exon flanked by two small introns; as translation of this sequence results in a truncated protein, these variants were also classed as non-functional (i.e., with the unspliced variants).

### Transposition rate estimation—fly crosses

Dysgenic and non-dysgenic F1 males were produced by reciprocal Cro18 x attached-X crosses at 28.5°C, and then crossed to attached-X virgin females at 25°C. Dysgenic and non-dysgenic F1 females were produced by reciprocal Cro18 x M252 crosses at 28.5°C, and then crossed to M252 males at 25°C. We performed a total of 40 mass crosses (10 per cross type, each with 5-10 females and 5-10 males), turned over into fresh food vials every 2-3 days. DNA was extracted from F2 adults (males and females) that emerged from these crosses.

### Transposition rate estimation—DNA extraction and pooling strategy

We performed 2128 individual fly DNA extractions using one of the following kits and following manufacturer’s instructions: GeneJet Genomic DNA Purification Kit (Thermo Fisher Scientific, Waltham, MA, USA), Qiagen DNeasy 96 Blood & Tissue Kit (Qiagen, Hilden, Germany), Zymo *Quick*-DNA 96 Plus Kit (Zymo, Irvine, CA, USA) and Zymo *Quick-*DNA 96 Kit. We then pooled these extractions into 286 groups of ∼8 flies of the same type (Supplementary Table 1). Each pool was quantified with Qubit (using the Qubit dsDNA HS Assay Kit: Invitrogen, Waltham, MA, USA), and 1-2µg of DNA (whenever possible) were subjected to shearing on a Bioruptor Pico sonicator (Diagenode, Denville, NJ, USA) to an average size of 200bp. We then cleaned up and concentrated the samples using AMPure beads at 1.8X (Beckman Coulter, Brea, CA, USA) prior to library preparation.

### Transposition rate estimation—Library preparation

The library preparation strategy relies on PCR to target regions near the breakpoints of TE insertions to identify insertion locations, in a modified version of previously developed method (Grech et al. 2019). Library preparation was performed using the KAPA Hyper Prep Kit (Roche, Basel, Switzerland), with an AMPure bead clean-up step (1.8X) after each step. Library preparation steps were as shown in Supplementary Figure 1; linker and primer sequences are given in Supplementary Table 2.

To prepare the resulting 12 libraries (six for *roo*, used as a control, and six for *P-element*) for sequencing, we performed a final AMPure bead clean-up step at 0.9X to remove small fragments. They were subsequently quantified (using Qubit), and their traces were checked on a Bioanalyzer (Agilent), prior to equimolar pooling and sequencing. Sequencing was performed at the CGR (Liverpool) on one lane of an S4 flow cell on the Illumina NovaSeq 6000 (paired-end, 2×150bp), aiming for ∼2% of lane data recovery (40M reads or ∼12Gb of sequencing data).

### Transposition rate estimation—bioinformatic analysis

Pooled libraries were de-multiplexed and trimmed for the presence of Illumina adapter sequences using Cutadapt v1.2.1, with parameter -O 3 (Martin 2011). Reads were further trimmed using Sickle version 1.2 (Joshi and Fass, 2011), with a minimum window quality score of 20. Trimmed reads shorter than 15bp were discarded. The resulting data were then further processed using custom python scripts to identify individual pools of eight flies, trim primer sequences, and remove PCR duplicates. Primer sequences were trimmed using cutadapt v4.5 with parameters --minimum-length 25 --action=trim --discard-untrimmed. PCR duplicates were identified from the random 4-mers in the primer sequences and by sequence similarity (Levenshtein distance < 10).

The trimmed, demultiplexed, and deduplicated reads were then mapped to specially prepared reference genomes with bwa mem v. 0.7.17. For the P-element libraries, a P-element consensus sequence was concatenated to the Dsim3.1 reference genome (which lacks P-element sequence). For *roo* libraries, the reference genome was prepared by mapping the *roo* consensus sequence to the Dsim3.1 reference with minimap -x asm20, masking the mapped regions in the reference using the maskfasta command in bedtools v 2.31.0 (Quinlan and Hall 2010), and then concatenating the *roo* sequence.

A custom python script was used to filter for reads mapping to the expected location in the TE sequence (immediately adjacent to the TE primer), alongside their read pair. We then used samtools depth (v.1.18) to calculate the depth at each position. Regions covered by at least 3 reads in a sample were retained for further analysis.

To identify insertions that were unique (and thus potentially represent new insertions), we intersected across samples with bedtools multiinter, and merged adjacent regions within 1kb with bedtools merge. The resulting tab delimited files were analysed using a custom R script to find regions unique to a library, which were considered potential new transposition sites. All custom scripts used in the transposition analysis are available at https://github.com/andrea-betancourt/transposition.

To analyse putative new insertions for a bias toward origins of replication (ORIs) (Spradling et al. 2011), we first annotated ORIs in the *D. simulans* Prin3.1 reference based on sequence homology with evidence-based annotations of ORIs in *D. melanogaster*, following (Spradling et al. 2011; Kofler et al. 2015) and (Langmüller et al. 2023). Briefly, we selected ORIs from the Flybase annotation (https://flybase.org; dmel-all-r6.65.gff), extracted the sequence with bedtools v. 2.31.1 getfasta, and mapped this sequence to the *D. simulans* reference using minimap2 v. 2.28 (Li 2018) using parameter -x asm20. Alignments were filtered for mapping quality 30, and lengths between 100-2000 to eliminate short matches and very long matches that preclude precise localisation. We used bedtools intersect to count putative new insertions (using both dysgenic and reciprocal insertions) overlapping with the origin annotation by at least one nucleotide.

## Results

We tested for P-element activity and dysgenesis-associated phenotypes in *D. simulans* males obtained from reciprocal crosses between M-type lines (lacking P-elements) and P-type lines (containing P-elements) at 28.5°C. In the direction of the cross with an M-type maternal line, which lacks maternal suppression of the P-element, offspring are expected to suffer from P-element activity resulting in dysgenic phenotypes—we refer to these offspring as dysgenic. The offspring of the reciprocal cross benefits from maternal P-element suppression— we refer to these as reciprocal controls. Normally, F1 males from reciprocal crosses differ in their X- and Y-chromosomes. To obtain F1 males with identical nuclear genotypes, we used an attached-X stock as the M-type, following Engels (1979; Figure 1). In the offspring of females from this stock, the usual inheritance of sex-chromosomes is reversed, with male offspring resulting from the fusion of Y-bearing oocytes with X-bearing sperm. In the reciprocal cross, sex chromosomes are inherited normally. As a result, reciprocal F1 dysgenic and control males have sex chromosomes that differ in the parent of origin, but which are genetically identical, with Y-chromosomes from the attached-X stock, and X-chromosomes from the P-type stock.

### Elevated P-element expression in dysgenic flies

We first examined F1 males for evidence of P-element de-repression in the dysgenic vs. control males in expression of the P-element. We sequenced the transcriptomes of dissected testes tissue from 2-4 day-old (young) and 21-day-old (old) F1 males from each direction of the reciprocal crosses (library characteristics shown in Supplementary Table 3). We analysed the effect of dysgenesis on TE expression, using a redundant library of *Drosophila* TEs (modified from Chakraborty et al., 2021). We found that only the P-element was differentially expressed at an FDR ≤ 0.05 between dysgenic and reciprocal males, with ∼3-fold higher expression in dysgenic testes (log2FC = 1.78, adjusted *p* = 0.02; Figure 2).

**Figure 2.**
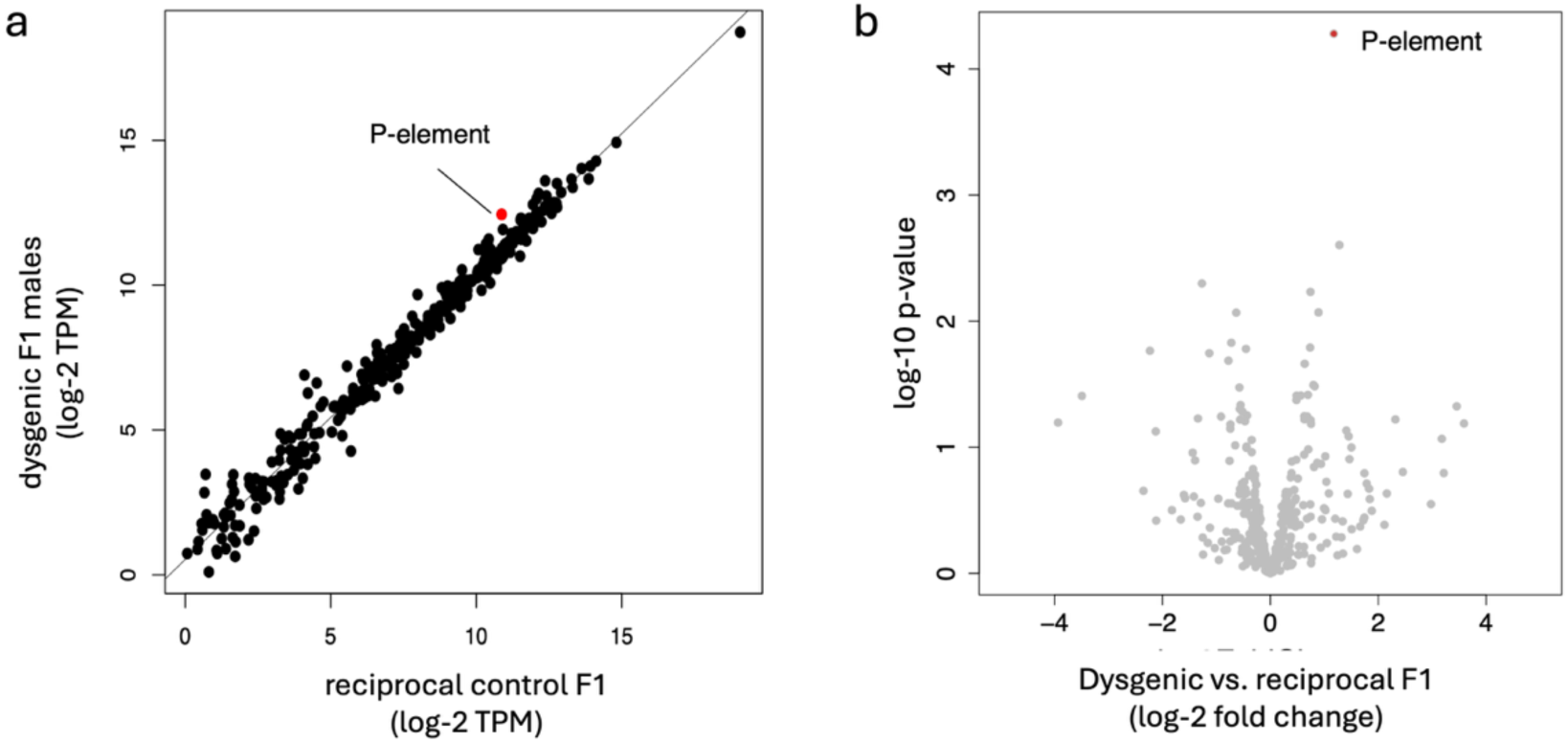
Expression of TE in testes from dysgenic vs. reciprocal males. A) Normalised expression of TEs from dysgenic and reciprocal flies, with the P-element highlighted in red. B) Volcano plot showing the comparison between testes samples from dysgenic and reciprocal flies. The log of the unadjusted p-value is shown on the y-axis; TEs with adjusted p-values < 0.05 and a log-2-fold-change > 1 are in red. For this analysis, young and old flies were binned together.

We also compared the F1 flies to the P-type parent—note that we used data only from young flies as we lack data for old parental testes, and that the F1 flies are expected to have roughly half the P-element content of the P-type parent. Nevertheless, P-element was elevated in F1 flies compared to the P-type stock, consistent with strong de-repression in dysgenic F1s (6.3-fold difference; Cro18 vs dysgenic: log2FC = −1.84, adjusted-*p* = 1.42e-17) and mild de-repression in reciprocal controls (1.8-fold difference; Cro18 vs reciprocal: log2FC = −0.580, adjusted-*p* = 0.0262; Supplementary Figure 2). The mildly elevated P-element expression in reciprocal F1 flies is consistent with reports of mild dysgenesis effects in this direction of the cross (Kidwell et al. 1977).

As there is some evidence that dysgenic *D. melanogaster* females regain P-element suppression as they age (Khurana et al. 2011), we compared subsets of the data from both young and old F1 males. Restricting the analysis to old males shows non-significant ∼2.1-fold higher expression in dysgenic vs. reciprocal males (log2FC = 1.11; adjusted *p* = 0.36). Restricting to young males, shows the P element had ∼2.3-fold higher in dysgenic males (log2FC of 1.2, adjusted *p* = 0.45; Supplementary Figure 2). That neither difference is significant is likely due to the low replication (*<u>n</u>* = 2) in the reduced data set, and overall, there was no evidence for a difference between young and old males in P-element expression.

Because suppression of splicing of the third intron of the P-element (IVS3) is thought to be a major regulator of P-element activity (Teixeira et al. 2017), we tested for a differences in the rate of canonical splicing of this intron between reciprocal F1s. Interestingly, dysgenic F1s often showed a retention of a mini exon of 32bp within IVS3, yielding a transcript encoding a non-functional transposase. Dysgenic F1s also had proportionally more fully spliced IVS3 transcripts resulting in transcripts encoding a functional transposase compared to controls (Supplementary Table 4; functional vs. nonfunctional transcripts: dysgenic = 32 vs 829; reciprocal 0 vs. 166; Fishers exact test, *p* = 0.0057). This suggests that dysgenic males not only have higher P-element expression, but that splicing of those transcripts is more efficient, resulting in more functional transposase expression than controls.

### P-element transposition

We next asked if the de-repression of the P-element seen in dysgenic F1 flies leads to an increase in germline transposition. We used targeted sequencing (using a method adapted from Grech *et al*, 2019) to compare P-element transposition in dysgenic flies vs. reciprocal controls. We sequenced P-element insertion breakpoints from the offspring of four types of flies: *i)* dysgenic and *ii)* reciprocal F1 males from an attached-X M x P cross, and *iii)* dysgenic and *iv)* reciprocal F1 females from a standard M x P cross. Each F1 was backcrossed to the M-type parent, and the offspring sequenced in pools of 8 flies, with a total of ∼500 backcross offspring for each type, *n* = 2096 flies in all. New P-element insertions in the F1 germline are expected to appear in heterozygous insertions in new locations in the backcross progeny. We thus counted P-elements as putative new insertions if they were unique to a pool (with a breakpoint > 1kb from other insertions in the library). To exclude PCR duplicates, we included only reads that contained unique sequences, including that of a random nucleotide tag (or Unique Molecular Identifier, see Materials and Methods).

In total, we recovered 666 of these putative new P-element insertions from the sequenced backcross progeny of the four types of F1 flies (Figure 3A). Dysgenic flies of both sexes had a significantly higher transposition rate than reciprocal controls, with ∼ 7-fold more putative new insertions (GLM with quasiPoisson errors, new transposition count ∼ sex of parent + cross direction (HD|R) and offset = number of flies in the pool; coefficient for HD vs. R = 1.927 on a log-scale, or a ∼6.9-fold difference; *χ^2^* = 155.2, df = 1, *p* < 2e-16). Dysgenic F1 females had a marginally higher transposition rate than males, 1.3-fold (GLM as above, *χ^2^* = 6.467, df = 1, *p* = 0.011). However, the effect of sex in this assay cannot be distinguished from genetic background, as different crosses were used to produce F1 males and females. Note that this difference cannot be explained by somatic transposition in dysgenic and non-dysgenic backcrosses used to produce the sequenced flies (Supplementary Table 1; 1.6 fold higher rate for the non-dysgenic cross; quasi-Poisson GLM restricted to data from F1 reciprocal flies only, new transposition count ∼ cross type; *χ^2^* = 2.468 df = 1, *p =* 0.116).

**Figure 3.**
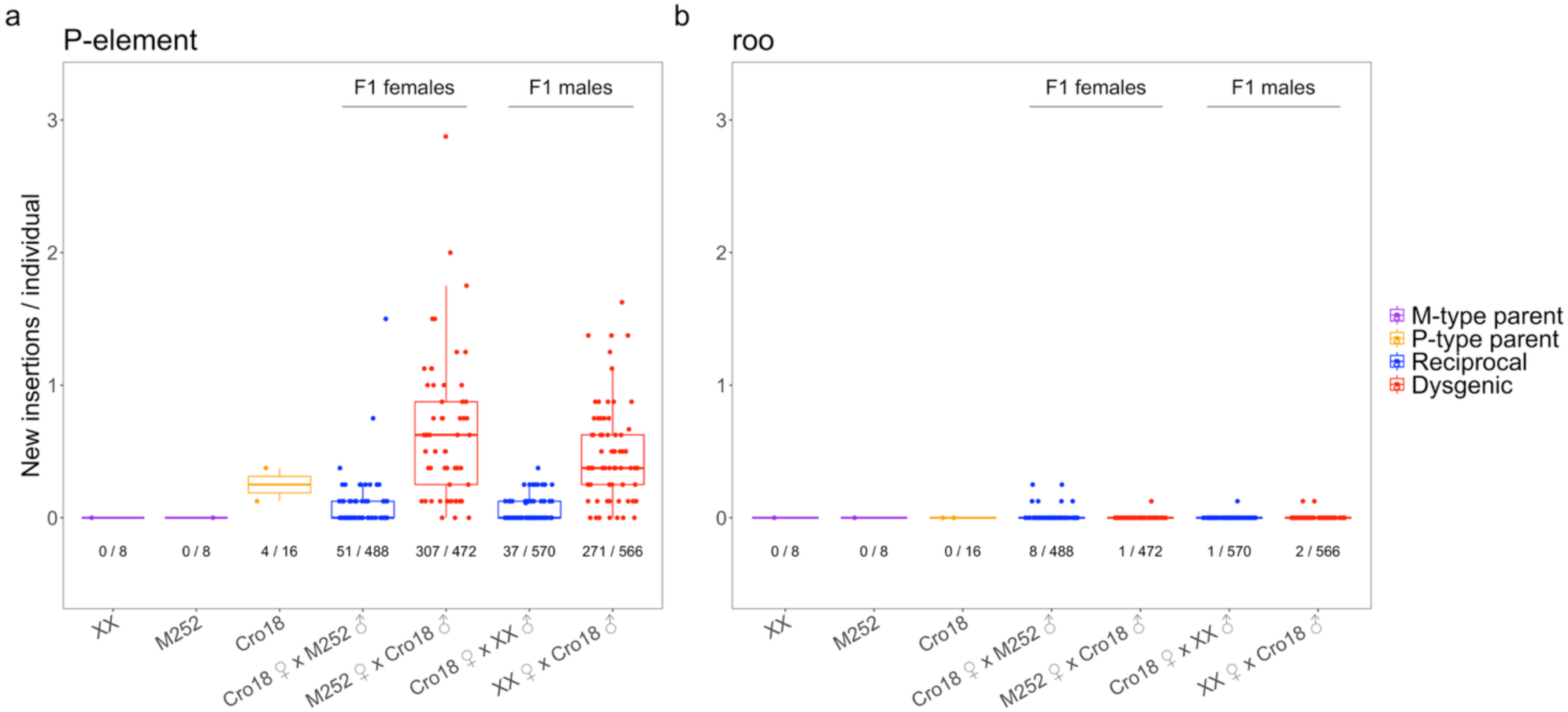
Transposition in dysgenic and reciprocal flies of both sexes. Putative new insertions occurring in F1 individuals were obtained by targeted sequencing of backcross offspring, as described in the main text. Boxplots and points show insertions recovered only in one sample, and which thus may correspond to new insertions in F1 females and males of A) the P-element, and B) roo. Numbers under the boxes are numbers of new insertions over total number of backcross progeny sequenced, and each point represents the insertions recovered from a library of ∼8 pooled flies, divided by the number of flies in that pool.

As a control, we performed the same type of assay for the retrotransposon *roo*, one of the most common transposable elements in *D. simulans* (Biemont et al. 1999; Kofler et al. 2015), using the same pools of flies as input for the *roo* assay. We identified only 12 putative new *roo* insertions, with most of these (*n* =8) occurring in the reciprocal F1 females; a marginally non-significant increase (Figure 3B; quasi-Poisson GLM *χ^2^* = 3.63 df = 1, *p =* 0.056).

We estimated 8-13 copies of P-element on average in the F1 offspring of the Cro18 strain, and that 56.5% of dysgenic backcross progeny had new insertions, implying a transposition rate of 0.0046 – 0.091 per individual per P-element copy. This estimate is high, but within the range of unrepressed transposition rates of 0.001 to 0.1 estimated in *D. melanogaster* (new insertions/element/genome; reviewed in Kelleher (2016). P-elements have an apparent insertion preference for origins of replication, or ORIs (Spradling et al. 2011), with natural and experimental populations both showing an enrichment of insertions in these locations (Kofler et al. 2015; Langmüller et al. 2023). We analysed these putative new insertions for a similar bias, and found a 15-fold enrichment, with 20.5% of the insertions overlapping with our *D. simulans* ORI annotations, which constituted roughly 1.35% of the reference genome (χ^2^ = 1890.4, df = 1, *p* < 2.2e-16).

### Transcriptional phenotype of dysgenic testes and germ cell underrepresentation

We next compared the transcriptional phenotypes of dysgenic males and reciprocal controls using analysis of RNAseq data. A principal component analysis (PCA) separated parental strains on PC1 (with F1s intermediate between them; 50% of the variance) and roughly separated young and old flies on PC2 (15% of the variance; Supplementary Figure 3). We first analysed the effect of dysgenesis on global gene regulation. We found 1,873 genes that were at least 2-fold significantly differentially expressed (DEGs; FDR ≤ 0.05) between dysgenic vs. reciprocal flies, with 733 up-, and 1,140 down-regulated in dysgenic flies (Figure 4A; Supplementary Figure 4).

**Figure 4.**
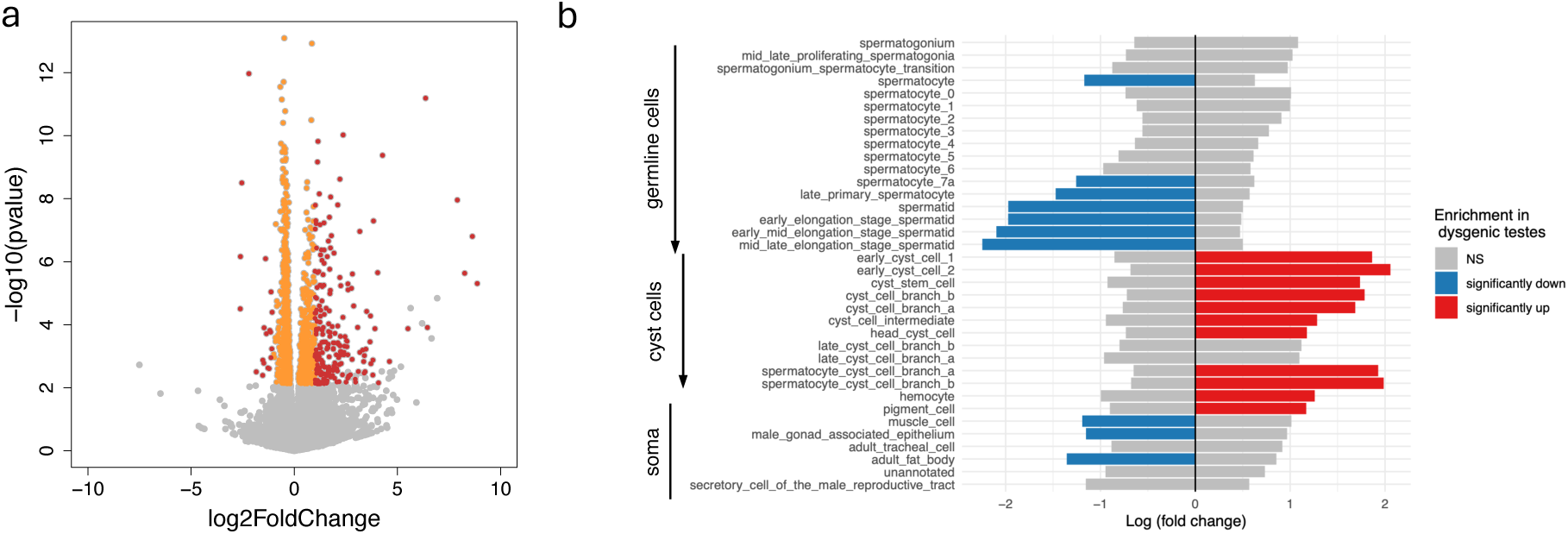
**A)** Effect of dysgenesis on global gene regulation. The volcano plot shows comparison between testes samples from dysgenic and reciprocal flies. The y-axis shows the log of the unadjusted *p-value*; Genes with adjusted p-values < 0.05 are shown in orange, those with adjusted p-values < 0.05 and a log-2-fold-change > 1 are show in red. **B)** Analysis of cell type differences between dysgenic vs reciprocal testes. Log fold changes refer to fold-changes in cell types inferred from differential gene expression in bulk testes tissue. Cell types are based on Raz et al. (2023), and the order from top to bottom within germline and cyst cells categories corresponds roughly to the order of differentiation. Significance is based on *p*-values < 0.05 after adjustment for multiple testing, assessed by bootstrapping.

To shed light on the biological relevance of DEGs between dysgenic and reciprocal flies, we performed a gene ontology (GO) term over-representation analysis, focusing on the DEGs shared between the old and young age classes (Supplementary Figure 5; Supplementary Tables 5 and 6). Because programmed cell death occurs in mitotically dividing germline stem cells and cystoblasts of dysgenic *D. melanogaster* females (Tasnim and Kelleher 2018), we inspected GO-terms associated with cell death, DNA damage repair and apoptosis (‘apoptosis’, ‘programmed cell death’, ‘cell death’, ‘site of double-stranded break’, ‘double-stranded break repair’), but found no evidence that these genes were more highly expressed in dysgenic than non-dysgenic males. Specific genes associated with cell death and the DNA damage response, including *Chk2*, *p53* (Tasnim and Kelleher 2017), and *Myc* (Ota and Kobayashi 2020) were also not differentially expressed in dysgenic vs. reciprocal males (p53: log2FC = −0.10; adjusted *p* = 0.66, *Chk2*-ortholog, *Lok*: log2FC = 0.31; adjusted *p* = 0.09, and *Myc*: log2FC = 0.47; adjusted *p* = 0.41).

However, if apoptosis is transient in males as it is in females, the gene expression data and differential expression may represent an over- or under-representation of specific cell types in the testes, rather than specific gene-level transcriptional responses. To investigate this possibility, we made use of available single-cell sequencing data from (Raz et al. 2023) to ask whether the shifts in expression among our genes might reflect a shift in cell-type composition. Analyses of these shows a significant depletion of several types of mature sperm cells, and an increase in somatic cyst cells, suggesting a loss of sperm cells at some point during spermatogenesis (Figure 4B). Consistent with this idea, the effect of cell death appears to compound with age: old males harbour more mature sperm than young males (Sanghvi et al. 2025) and dysgenesis appears to disrupt expression of more genes in old vs. young males (1,192 DEGs in old males vs. 501 in young males; Supplementary Figure 4).

### Consequences of P-element dysregulation in dysgenic males on fertility

The deleterious effects of P-element activity on fertility have been well-documented in *D. melanogaster* males and females (Kidwell et al. 1977; Engels and Preston 1979), with fertility less severely impacted in male *D. melanogaster* compared to females (Engels and Preston, 1978). The analysis above suggests that P-element may also cause fertility defects in male dysgenic flies by inducing the death of mature sperm cells. We tested reciprocal F1 males for fertility defects by measuring offspring production of 26 dysgenic and 30 reciprocal males mated to P-type females from the Cro18 stock. Dysgenic males showed more apparent sterility, with no offspring produced (*n =* 4 of 26 dysgenic males vs. 0 of 30 controls; Fisher’s exact test, *p* = 0.041). Dysgenic F1 males also showed a lower peak in offspring production than reciprocal controls (Figure 5A; day 18 mean ± SD 6.02 ± 4.83 for dysgenic vs 15.77 ± 9.96 for reciprocal; quasipoisson (link = ‘log’) glm *P* = 2.83e-05. However, there was no overall difference in the total offspring production (Figure 5B; mean ± SD 86.7 ± 60.2 for dysgenic vs 100.6 ± 47.6 for reciprocal, negative binomial glm *P* = 0.504).

**Figure 5.**
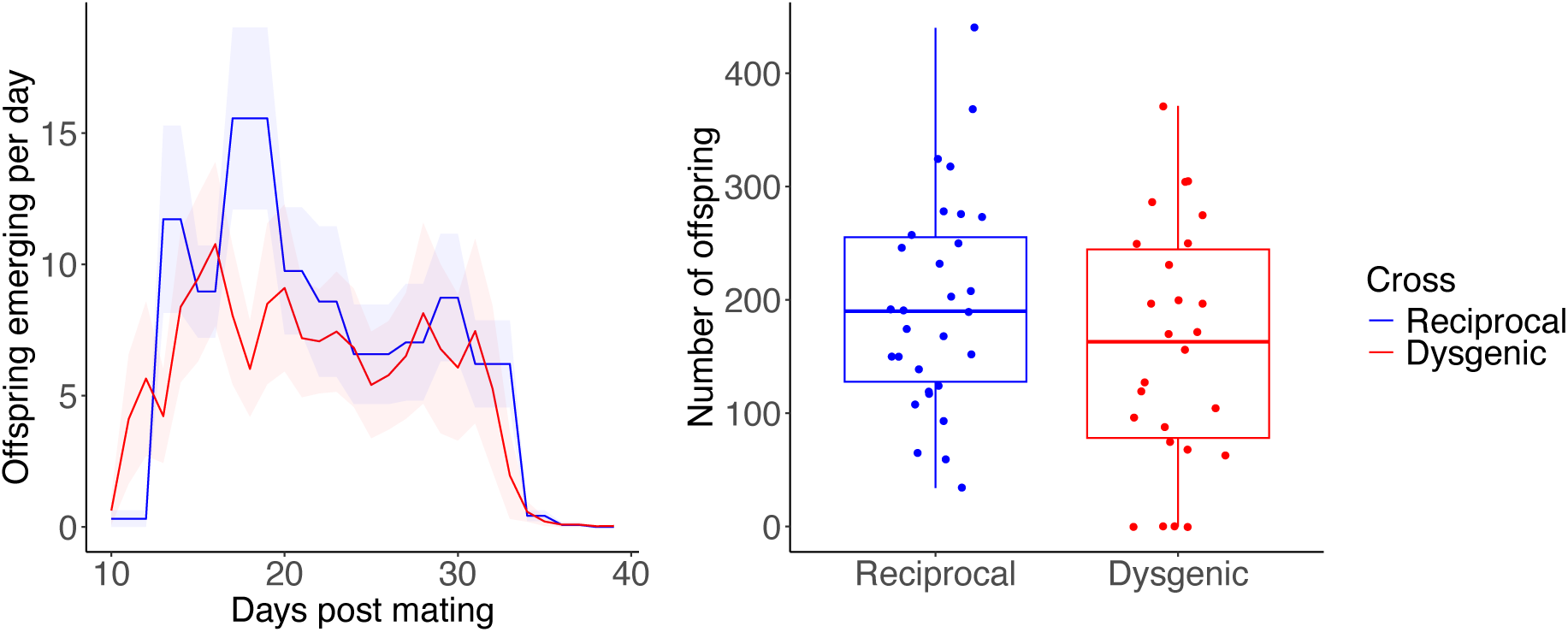
Offspring production from dysgenic and non-dysgenic males. **A.** The number of male and female offspring produced per day following parental mating. Offspring from dysgenic crosses are shown with a red line and offspring from non-dysgenic crosses are shown with a blue line. Ribbons indicate 95% confidence intervals. **B.** Average number of offspring produced from dysgenic and reciprocal males. The box plots display the upper and lower quartiles, the median and the range. Points represent each measurement obtained.

### Suppression of TE activity in the testes via the piRNA pathway

Small RNAs, especially piRNAs (small RNAs that associate with PIWI protein), are thought to be the main germline defence against transposable element activity in flies. In *Drosophila* oocytes most piRNAs are generated *via* the germline-specific ping-pong amplification loop, wherein *Aub* and *Ago3* cooperate to generate sense and anti-sense piRNAs from cognate TE mRNAs (reviewed in Czech et al., 2013; Senti et al., 2015). Though less studied in testes, the ping-pong pathway is also active in the male germline (Aravin et al. 2004; Vagin et al. 2006; Nishida et al. 2007; Saint-Leandre et al. 2020; Chen et al. 2021).

We assessed expression of important piRNA pathway genes in our testes transcriptomic data. We curated a list of 34 genes described as having important role in the piRNA pathway. Consistent with other work showing expression of piRNA-mediated silencing in male as well as female *D. melanogaster* (Saint-Leandre et al. 2020; Chen et al. 2021), all of the piRNA pathway genes are expressed, except for one, *Veneno* (had transcripts per million (TPM) > 1; Figure 6). Genes involved in the germline branch of the piRNA pathway in females, i.e., involving post-transcriptional silencing *via* ping-pong amplification (Brennecke 2007), such as *Argonaute-3*, *Aubergine*, *Piwi*, *Zucchini*, and *Vasa*, are also expressed.

**Figure 6.**
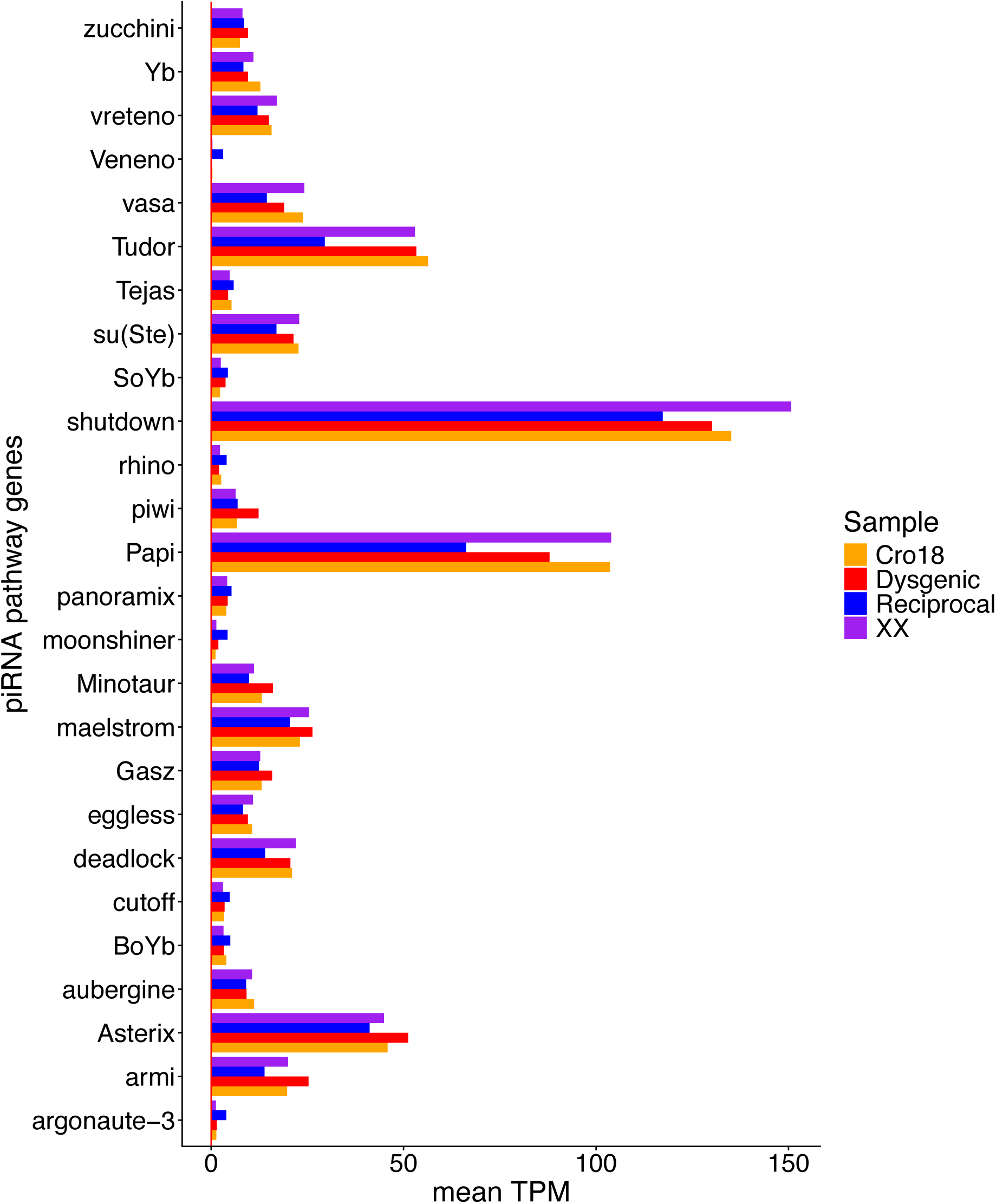
Expression of selected piRNA pathway genes in testes. Expression values are TPM averaged across replicates; the red vertical line corresponds to a TPM value of 1.

We also examined piRNA production in the testes by isolating small RNAs from young and old dysgenic and reciprocal testes and their attached X and Cro18 parents. TE-derived piRNAs are homologous to transposons and silence active transposons by cleaving their mRNAs (reviewed in Senti & Brennecke (2010). After filtering the small RNA data for read lengths between 23-32 bp, we recovered 0.4-39M reads from dysgenic and reciprocal young and old samples (Supplementary Table 7). Consistent with previous work and the analysis above, these data provide evidence that the ping-pong pathway is active in testes, despite the marginal expression of *Ago-3*, as our samples show the characteristic ping-pong signature of sense and anti-sense piRNAs that overlap by 10 nucleotides (Supplementary Figure 6A). Investigating piRNAs cognate to the P-element specifically, we see that Cro18 parents show moderate expression of these piRNAs (TPM=1278.61) as expected, while the attached-X parents essentially lack them, again, as expected (read counts of 0 and 2 per replicate, with the 2 reads possibly due to mild cross-contamination between samples). P-element cognate piRNAs also generally show a ping-pong signature (Supplementary Figure 6B).

### Relationship between expression of piRNAs, TE mRNA and P-element induced dysgenesis

Overall, dysgenesis did not appear to result in global TE dysregulation in our transcriptomic data, with only the P-element differentially expressed between dysgenic flies and reciprocal controls (see above). Similarly, there was little evidence for changes in piRNA composition between dysgenic and reciprocal control testes, or between these flies and the parental lines, with no piRNAs significantly differentially expressed between dysgenic and reciprocal males (FDR < 0.05; Figure 7). Restricting these comparisons to old or young samples similarly resulted in only small numbers of differentially expressed TEs or piRNAs (Supplementary Figures 2 and 7). Interestingly, for both TE mRNA and piRNAs, principal components analysis shows that the main variation among genotypes is driven by differences in expression of P-element and Shellder, another TE suggested to have invaded *D. simulans* recently, slightly before the P-element (Supplementary Figure 8; Scarpa et al. 2024).

**Figure 7.**
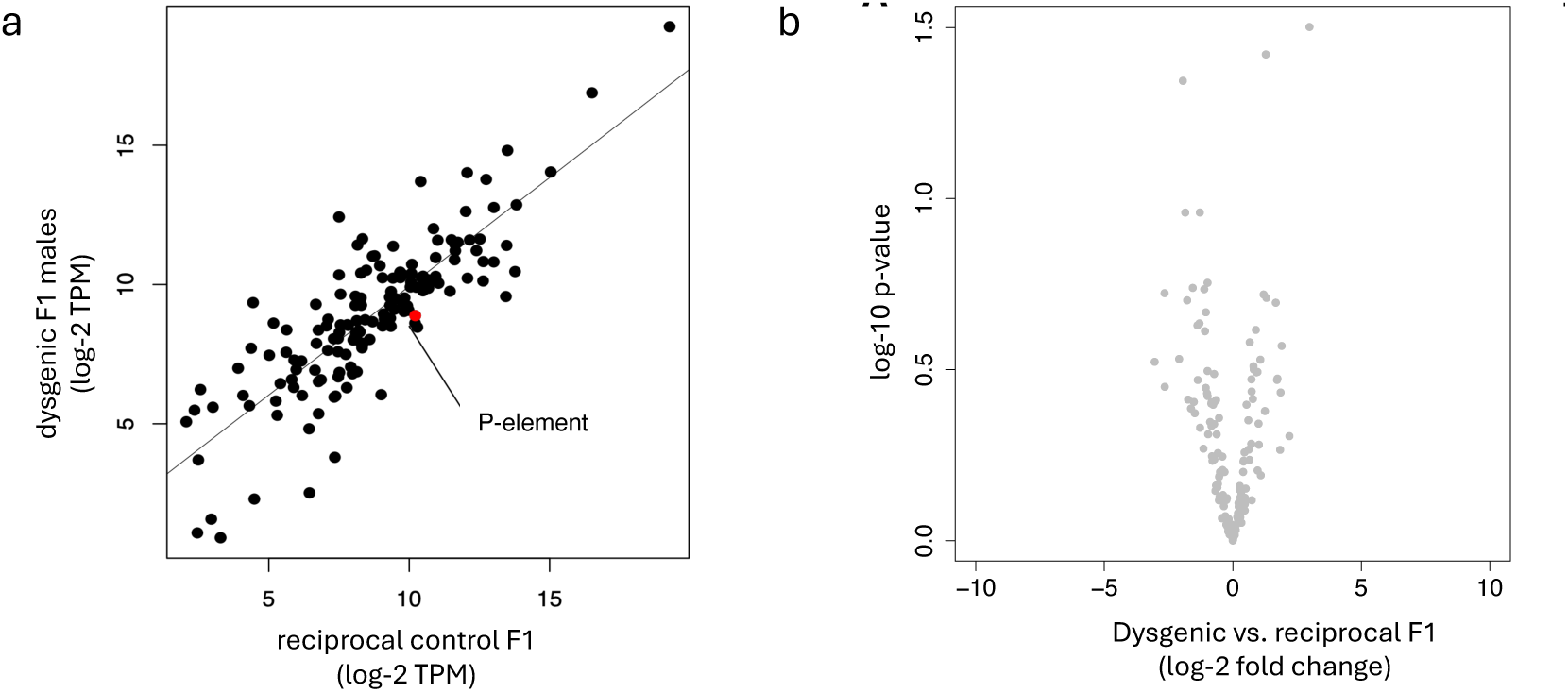
Expression of piRNAs expressed in testes from dysgenic vs. reciprocal males. A) Normalised expression of piRNAs from dysgenic and reciprocal flies, with the P-element highlighted in red. B) Volcano plot showing the comparison between testes samples from dysgenic and reciprocal flies. No TEs had adjusted p-values < 0.05 and a log-2-fold-change > 1.

To evaluate the signature of P-element dysregulation in dysgenic flies, we investigated the relationship between TE mRNA and their cognate piRNAs, both across all TEs and specifically in the P-element. For most TEs, the abundance of piRNAs and TE mRNAs is positively correlated, similar to results for ovarian piRNAs in *D. simulans* (Figure 8A; Lerat et al. 2017). This positive correlation is expected—highly expressed TEs will generate many piRNAs, as most piRNAs being secondarily derived from mRNAs during ping-pong biogenesis (Senti et al. 2015). We expected, however, that this relationship might change for poorly regulated TEs— i.e., they may have high levels of TE mRNA compared to piRNAs. Consistent with this idea, piRNAs are negative correlated with the levels of mRNA across samples, and show amongst the most negative relationship for any TE family (Figure 8B and 8C), and dysgenic individuals had the highest ratio of P-element mRNAs to piRNAs (excepting the M-type attached-X stock), mainly due to higher levels of mRNA (Supplementary Figure 8C).

**Figure 8.**
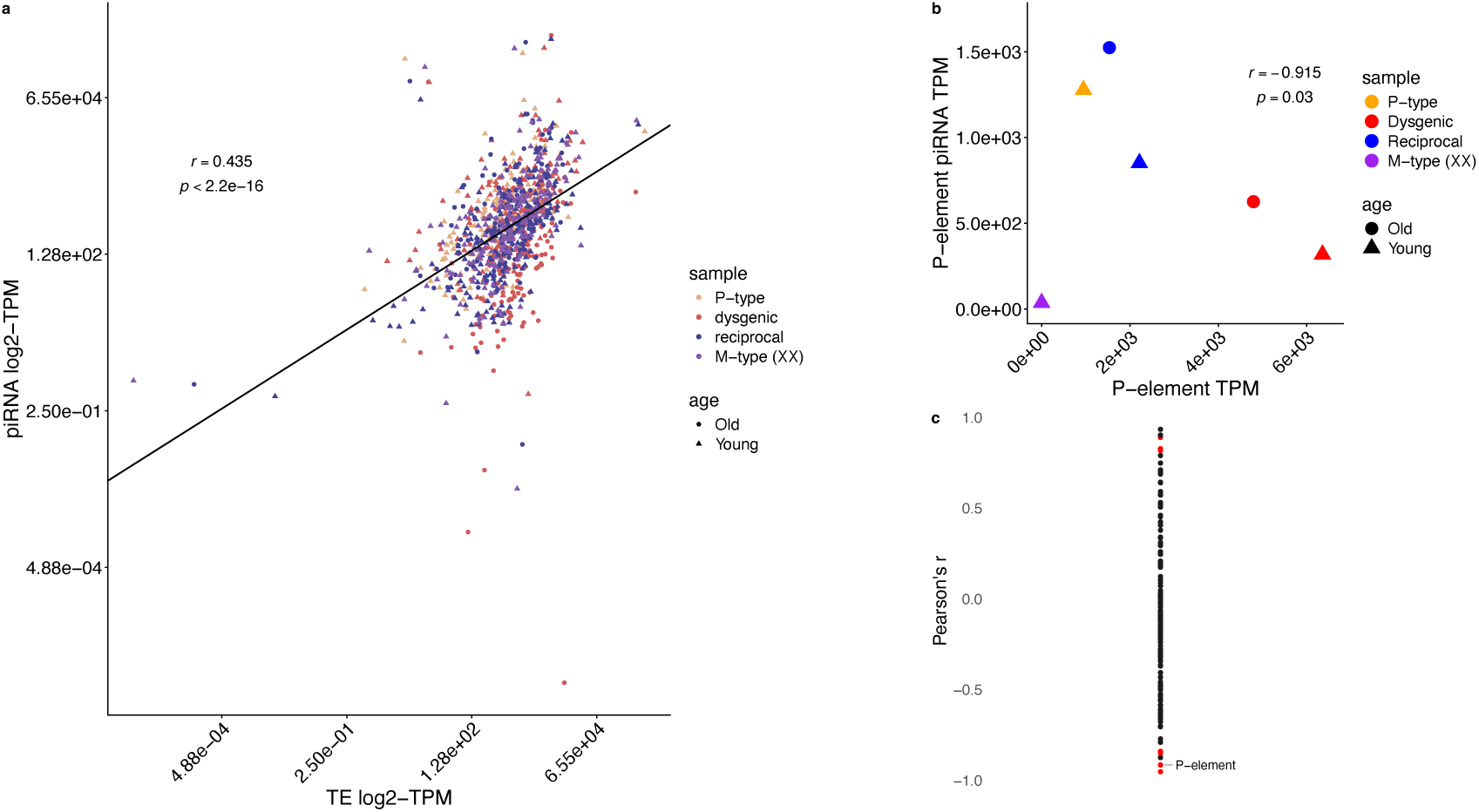
The relationship between piRNA and TE mRNA abundance measured as TPM values obtained from Salmon TE. A) Data for all TEs. A single line fit to all the data is shown for illustrative purposes. Analysis with linear models shows that the slope does not differ significantly for different samples; analysis with linear models shows that the slope differs significantly but slightly for different samples, with the slope varying from 0.445 to 0.650, B) data from the P-element. P-element data are shown on a linear scale to show the data for the M-type stock. C) data from all TEs expressed in at least three samples, showing the Pearson’s correlation coefficients for each family independently (for families occuring in at least three samples). Those with adjusted Bejamini-Hochberg adjusted *p* ≤ 0.05 are shown in red. To assess statistical relationships, we used a Pearson correlation tests on log₂(piRNA-TPM) and log₂(RNA-TPM) excluding values of zero.

## Discussion

Here, we examine P-element induced dysgensis in *D. simulans* males, and find that dysgenic males suffer P-element de-repression, high rates of P-element transposition, and the potential loss of mature sperm cells as inferred from patterns of differential gene expression in testes. One caveat, however, is due to the use of the attached-X stock. These crosses are a standard method by which *Drosophila* sex-chromosome inheritance is manipulated (Morgan 1922; Courret et al. 2019; Brown et al. 2020), and allowed us to control for nuclear genotype while manipulating the maternal cytotype. However, in these crosses, the paternal origin of the X and Y chromosomes differs between F1 males, and epigenetic effects due to the parent-of-origin cannot be disentangled from the effect of cytotype. Unlike in mammals, parent-of-origin effects are not expected to be common, and genomic imprinting appears absent, at least in females (Coolon et al. 2012). However, large-scale gene expression differences between F1 males in attached-X crosses in *D. melanogaster* have been attributed to such parent-of-origin effects (Lemos et al. 2014). We argue, however, that P-element activity, followed by DNA damage and apoptosis of germ cells offers a simpler explanation for the gene expression changes seen here than parent-of-origin effects. That is, if P-element activity results in apoptosis in males during mitotic proliferation of germ cells (as it does in females; Tasnim and Kelleher 2017; Lama et al. 2022), then post-spermatocyte sperm cells will be underrepresented in dysgenic testes, resulting in apparent down-regulation of germline expressed genes, and corresponding upregulation of somatically expressed ones.

We also see expression of piRNAs and piRNA pathway genes in *Drosophila* testes, consistent with previous work (Aravin et al. 2004; Vagin et al. 2006; Nishida et al. 2007; Saint-Leandre et al. 2020; Chen et al. 2021), though perhaps to a lower extent than in females (*cf.* Saint-Leandre et al., 2020). Some relevant genes are expressed at low levels in bulk testes transcriptomes likely due to a narrow expression niche—*e.g.,* the protein complex formed of Rhino, Deadlock and Cutoff that drives transcription of piRNAs in oocytes also forms a protein complex in testes, but restricted to early spermatogenesis (Mohn et al. 2014; Andersen et al. 2017; Chen et al. 2021). However, the higher expression of *Aub* relative to its partner in the ping-pong cycle, *Ago-3*, may be noteworthy. Proteins encoded by these two genes cooperate to effect ping-pong cleavage, binding anti-sense and sense TE mRNAs respectively (Aravin et al. 2007; Li et al. 2009; Nagao et al. 2010), and thus naively might be expected to occur in equimolar proportions at the protein level. Here, we see a mean 6.4-fold bias toward *Aub* mRNA, more extreme than the 2-fold bias seen in ovary-cell lines for *D. melanogaster* (Lau et al. 2009), but a clear ping-pong signature in testes piRNAs. Assuming that relative mRNA levels reflect relative protein abundance, this bias toward *Aub* mRNA is consistent with ping-pong cleavage in males being at least partly homotypic, involving only *Aub*, as has been previously suggested (Quénerch’du et al., 2016), and as seen in females where *Ago-3* has been knocked-down (Senti et al., 2015; Zhang et al., 2011).

The increased expression of P-element in testes of dysgenic males and absence of maternal cognate piRNAs might be expected to result in elevated P-element activity in dysgenic males compared to reciprocal controls. Indeed, we estimate higher transposition activity in dysgenic flies, consistent with what has been previously seen for *D. melanogaster* (Dowsett 1983; Biémont et al. 1990). In common with *D. melanogaster* with active P-elements, these insertions show a bias towards replication initiation origins, suggested to increase the efficiency of copy number increase (Spradling et al. 2011), or to escape purifying selection (Langmüller et al. 2023), indicating that at least some of the enrichment seen in natural populations (Kofler et al. 2015) is due to insertion bias.

There was little evidence of sexual dimorphism in transposition activity here, with comparable rates of new P-element insertions originating in either males or females, though note here sex is confounded with genetic background. However, there were milder consequences for this activity in males compared to females, consistent with work in *D. melanogaster* (Kidwell 1977). Even at temperatures where dysgenesis phenotypes are mildly expressed (25°C), *D. simulans* females showed high rates of sterility (∼50%) and low offspring production (∼40% of that of controls; Hill et al. 2016). Yet here, at a high temperature that induces extreme dysgenic phenotypes in females, males showed only moderate rates of sterility of ∼10%, and no overall difference in offspring production. These relatively mild fertility effects, coupled with similar transposition rates in males and females and potentially lower piRNA activity (Saint-Leandre et al. 2020) suggest that the P-element may spread more easily through males than females.

## Data availability

Sequence data is deposited in the European Nucleotide Archive (project accession PRJEB101158). Fly stocks are available upon request, subject to viability.

## Acknowledgments

We are grateful to C. Montchamp-Moreau and C. Courret for sharing the attached-X stock. We thank M. Kelly, C Ramirez and R. Wong for technical assistance, and D Jeffares and staff at the Centre for Genomics Research Liverpool, particularly J. Kenny, for project design advice. We thank A. Ludwig and J. Brookfield for critical reading of the manuscript, A. Ludwig and S. Chamberyon for helpful discussion, and two anonymous reviewers for their insightful comments.

---

**Supplementary Figure 1.**
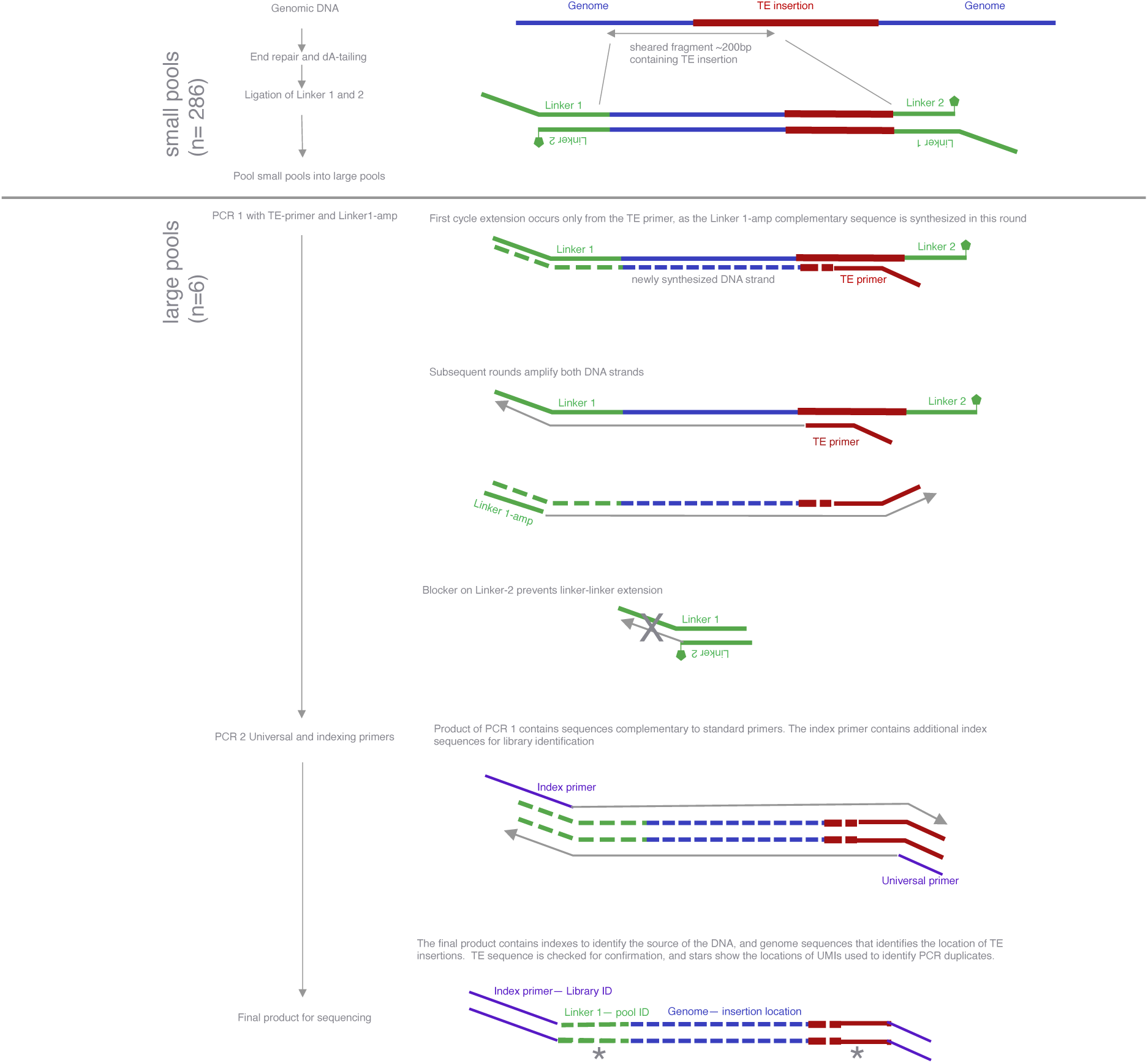
Library preparation protocol to sequence breakpoints of TE insertions, following Grech et al (2019). Linker and primer sequences are given in Supplementary Table 2. The initial input for library preparation was genomic DNA isolated from ∼8 flies and then pooled, as described in the Materials and Methods. *End-repair and A-tailing* was performed for each of the 286 pools of ∼eight flies. *Linker ligation* was also performed for each of the 286 pools of ∼eight flies. We ligated a forked linker, composed of Linker-1 and Linker-2. Linker-1 contains a 6nt barcode for pool identification, and a random 4-mer. Linker-2 contains a modified base on the 3’ end that prevents linker-linker amplification in the first PCR cycle (see below). *Pooling.* Following linker ligation, the barcodes in Linker-1 can be used to identify the specific original pool of ∼eight flies (“small pool”). Barcoded “small pools” were subsequently pooled into six “large pools”, such that each “large pool” is formed of ∼48 “small pools” (or approximately 8 x 48 = 384 flies). Subsequent PCR steps were performed on “large pools”. *PCR1.* In the first round of PCR, the ends of TEs and adjacent genomic regions were selectively amplified. Two primers were used, a TE-primer partly complementary to part of the TE of interest (P-element or roo), and Linker1-amp, a primer complementary to the unique part of Linker 1 (not present in Linker 2). In the initial cycle of this first PCR, extension occurred only from the TE primer (due to the 3’ blocker on Linker 2), which forms the binding site for the Linker1-amp. In subsequent cycles, both primers can then anneal and be extended, and both strands are amplified. *PCR2.* In the second round of PCR, standard Illumina primers (NEBNext Multiplex Oligos for Illumina) are used to create barcoded libraries for sequencing (each library corresponding to one “large pool” and TE target). This library preparation protocol was performed separately for P-element and roo, for a total of 12 sequenced libraries (6 ‘large pools’ for each TE).

**Supplementary Figure 2.**
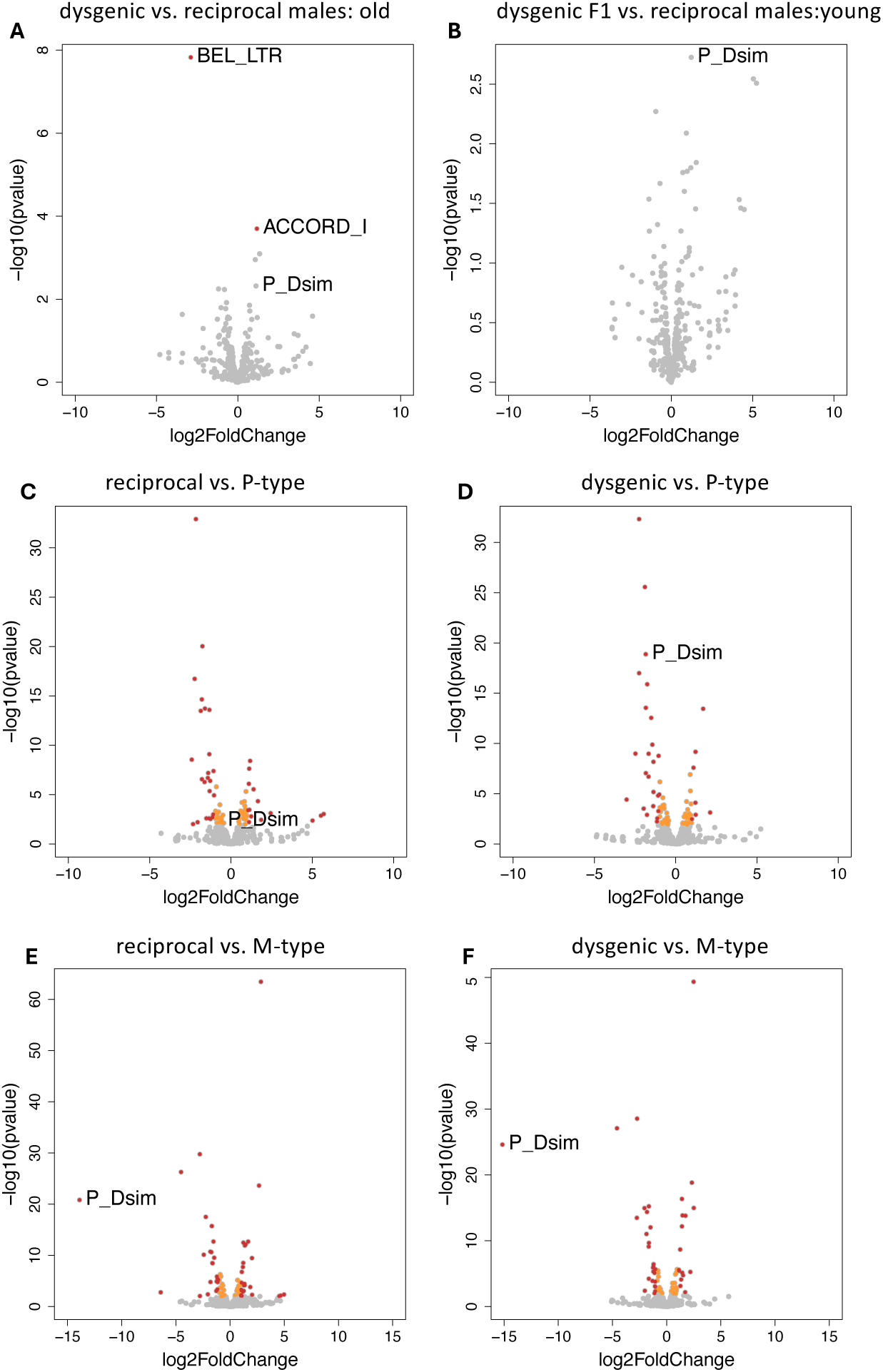
The effect of dysgenesis on TE expression in testes. Top row comparisons are between A) old, and B) young dysgenic F1 vs. reciprocal control males (BEL_LTR refers to long-terminal repeats of BEL, and not the full element). The middle row shows the P-type parent (Cro18) vs. C) reciprocal and D) dysgenic F1s, and the bottom row shows the M-type, attached-X parent vs. E) reciprocal and F) dysgenic F1s. TEs with adjusted p-values < 0.05 are shown in orange, those with adjusted p-values < 0.05 and a log-2-fold-change > 1 are show in red. The P-element is labelled regardless of significance.

**Supplementary Figure 3.**
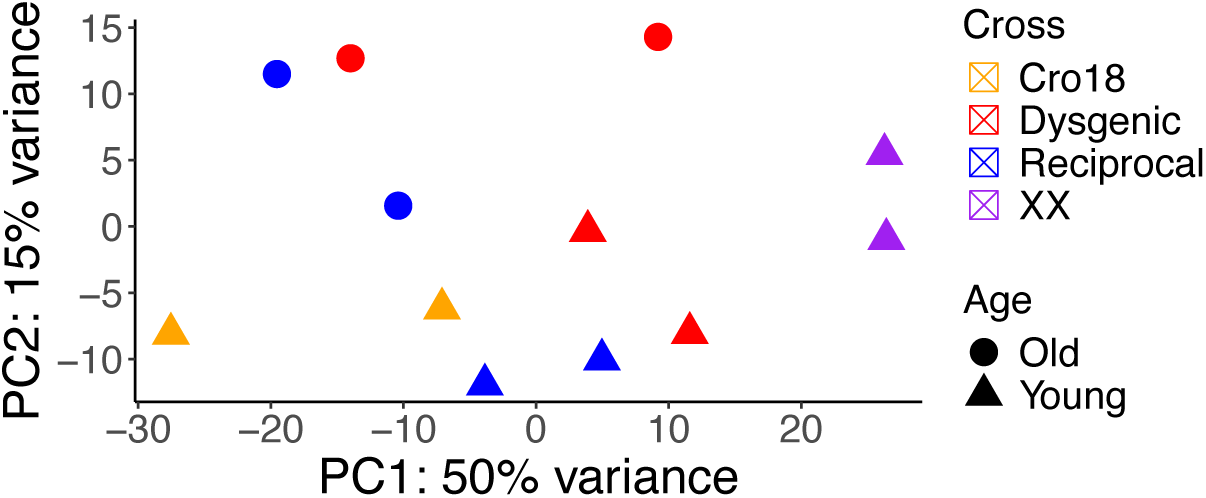
Principal component analysis coordinate plot showing normalised counts of transcripts from DESeq2, obtained from RNA-seq data quantified by Salmon.

**Supplementary Figure 4.**
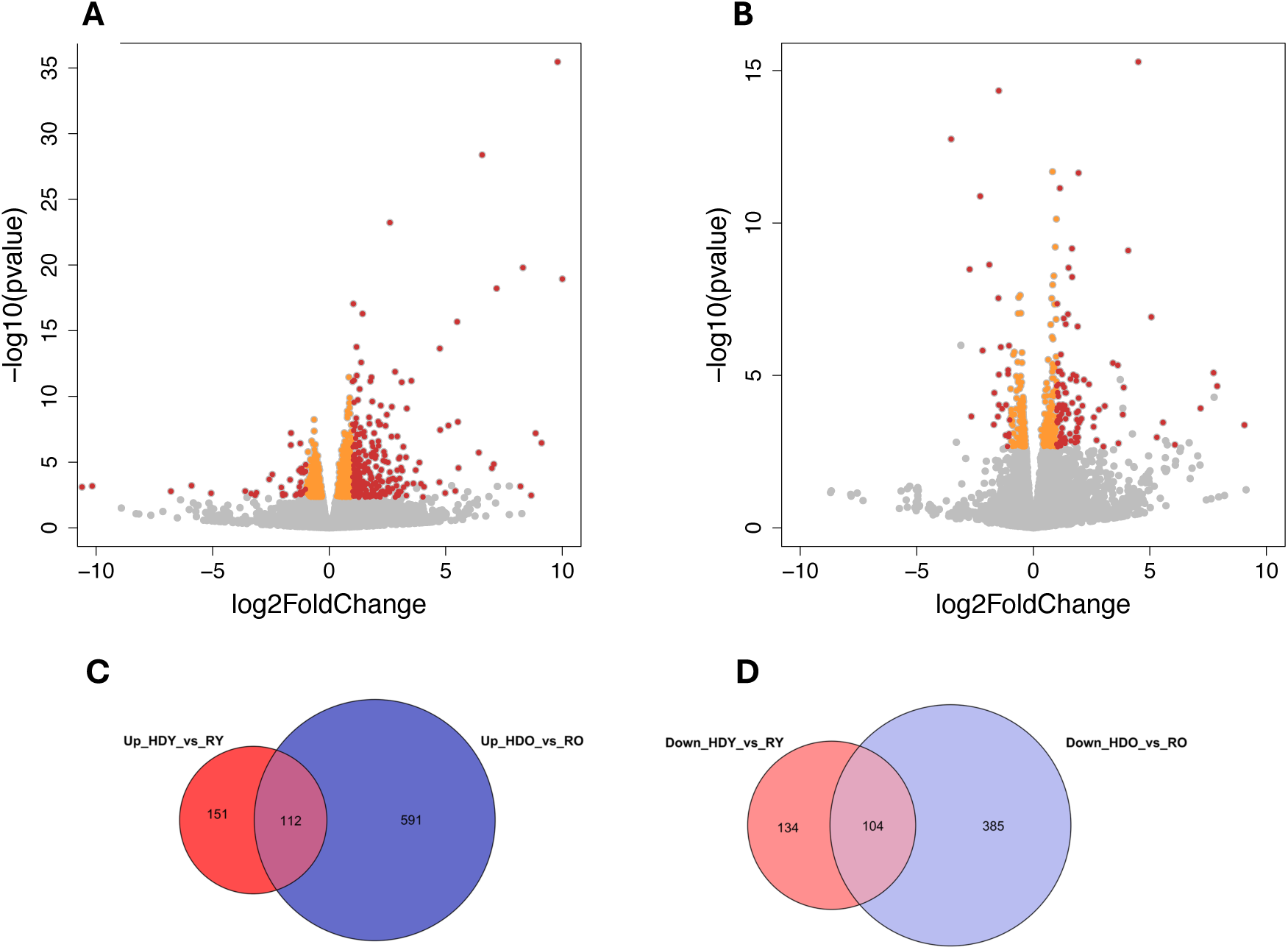
Effect of dysgenesis on global gene regulation in testes from young vs. old flies. Volcano plots show comparison between A) only old dysgenic and reciprocal flies, and B) only young dysgenic and reciprocal flies. Genes with adjusted p-values < 0.05 are shown in orange, those with adjusted p-values < 0.05 and a log-2-fold-change > 1 are show in red. The Venn diagrams show the number of genes with C) higher, and D) lower expression in dysgenic F1 vs. reciprocal controls, with comparisons done for the young and old age classes separately. Acronyms refer to hybrid dysgenic young (HDY), reciprocal young (RY) and hybrid dysgenic old (HDO) and reciprocal old (RO) samples. ‘Down’ and ‘up’ indicate whether the gene was down- or up-regulated in the first sample of the comparison (e.g., ‘Down_HDO vs RO’ indicates genes that were down-regulated in HDO compared to RO testes).

**Supplementary Figure 5.**
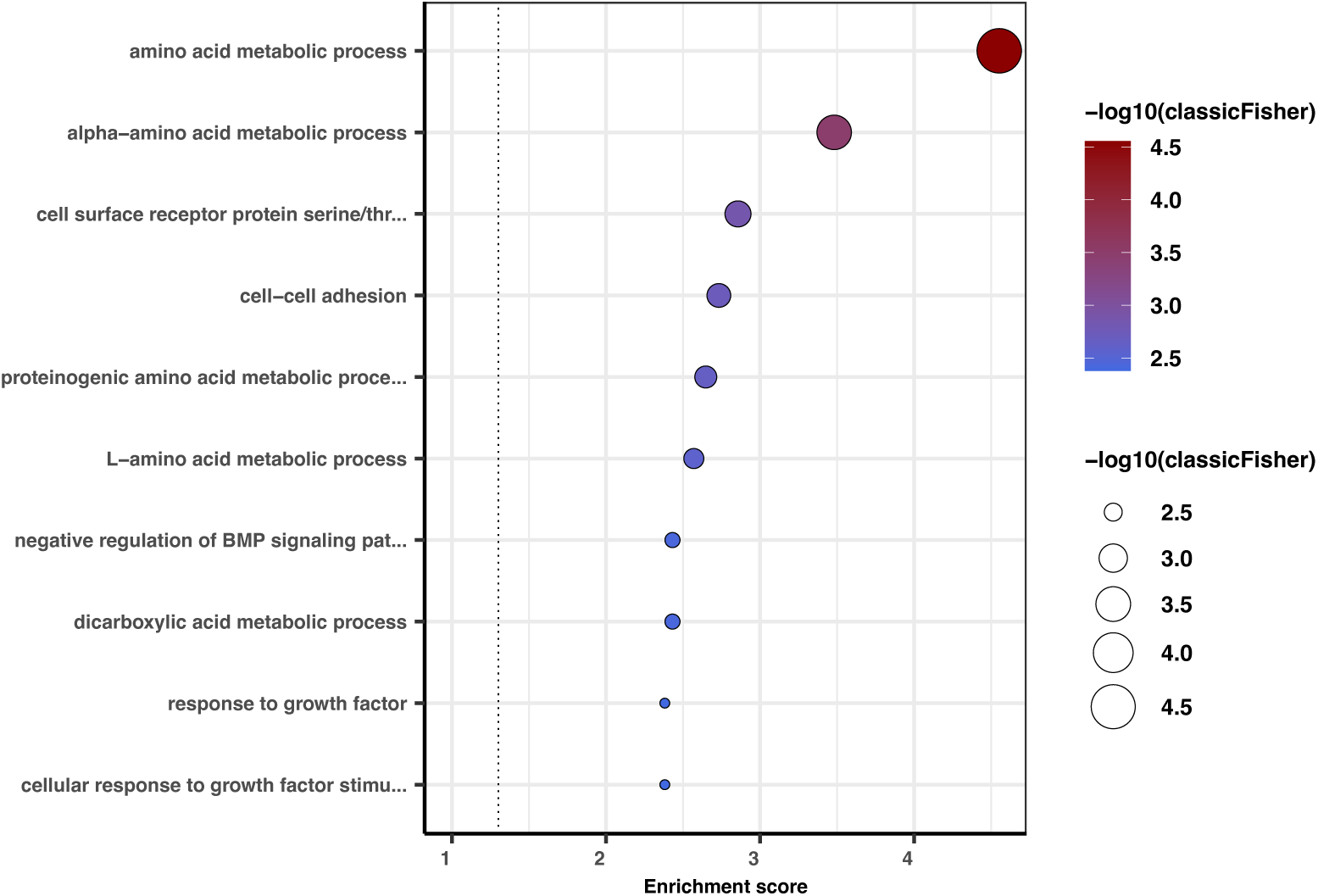
Gene ontology (GO) term over-representation analysis of genes differentially expressed between dysgenic and reciprocal testes shared among the old and young samples. The top 10 most significant GO terms are shown.

**Supplementary Figure 6.**
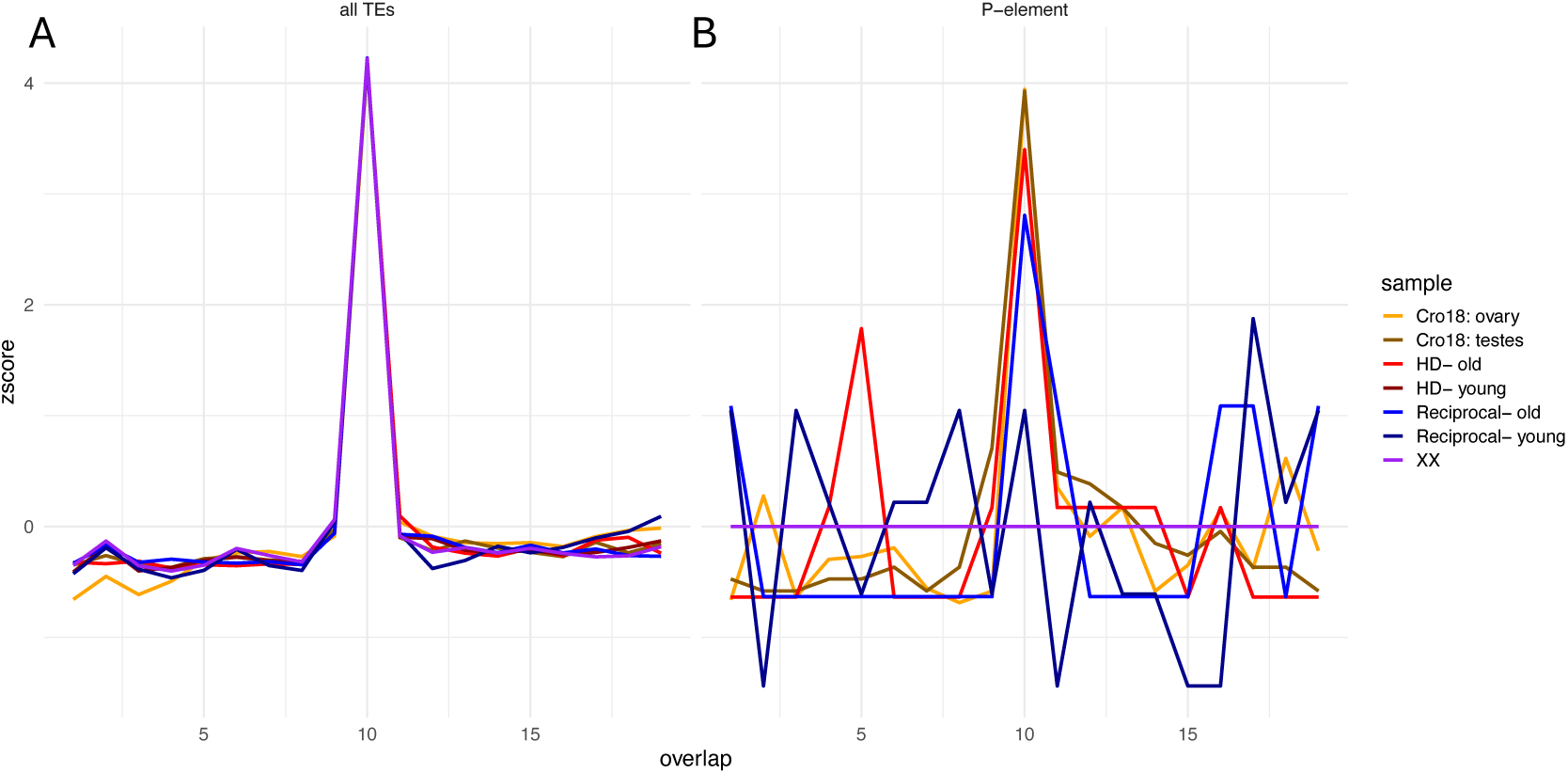
Ping pong signatures shown by an enrichment of sense and anti-sense piRNAs with a 10-bp overlap for piRNAs cognate to A) all TEs, and B) the P-element. Note that Z-scores > 2 indicate a significant enrichment at the alpha = 0.05 level. There were no overlapping P-element piRNAs in attached-X flies (XX), and few in the reciprocal young sample; all others show z-scores > 2 for overlaps of 10 nucleotides. Old dysgenic F1 vs. reciprocal males Young dysgenic F1 vs. reciprocal males

**Supplementary Figure 7.**
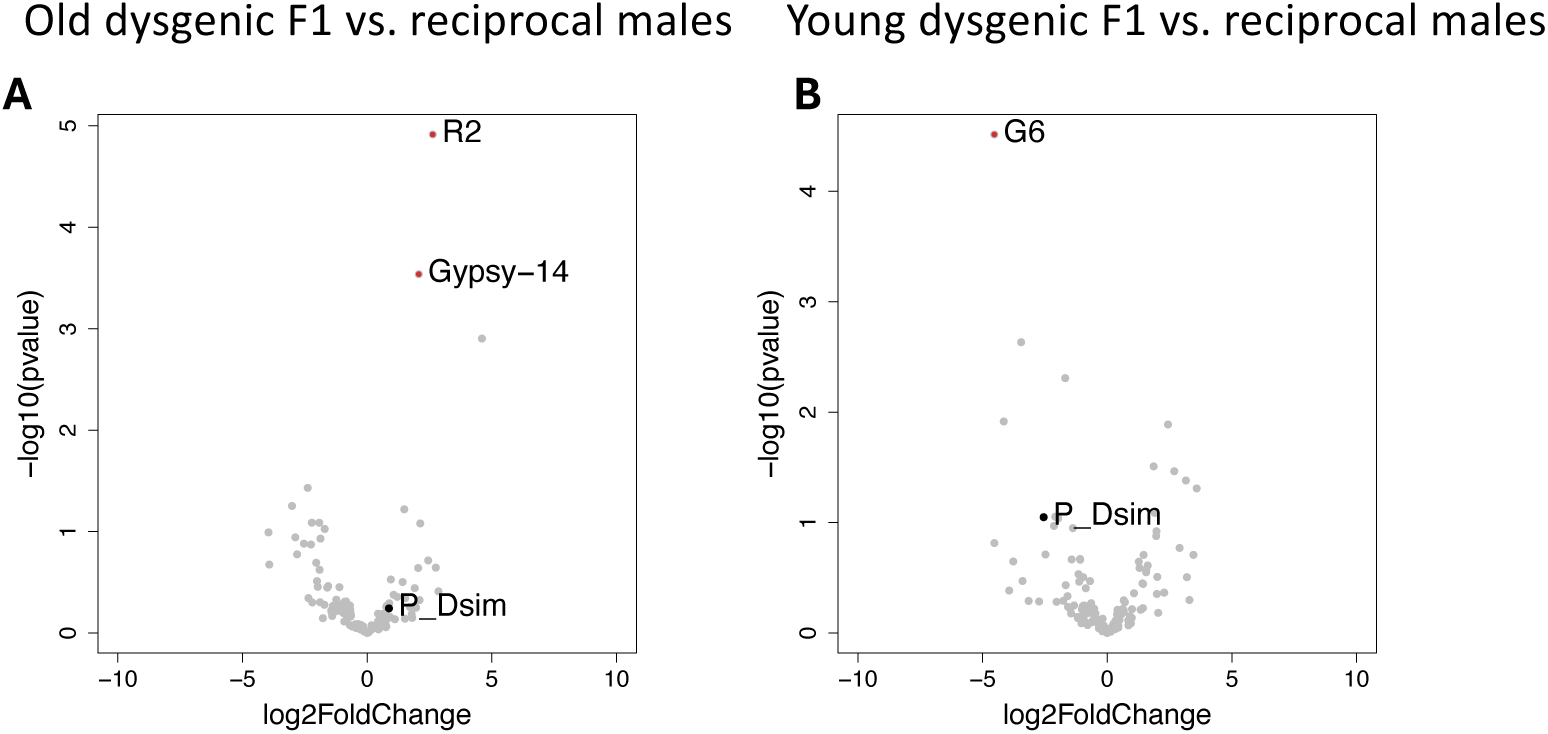
Expression of piRNAs expressed in testes from dysgenic vs. reciprocal males in different age classes. Shown are A) old testes dysgenic vs. reciprocal samples; and B) young testes dysgenic vs. reciprocal samples. Labels show names of cognate TEs; those with adjusted p-values < 0.05 and a log-2-fold-change > 1 are show in red.

**Supplementary Figure 8.**
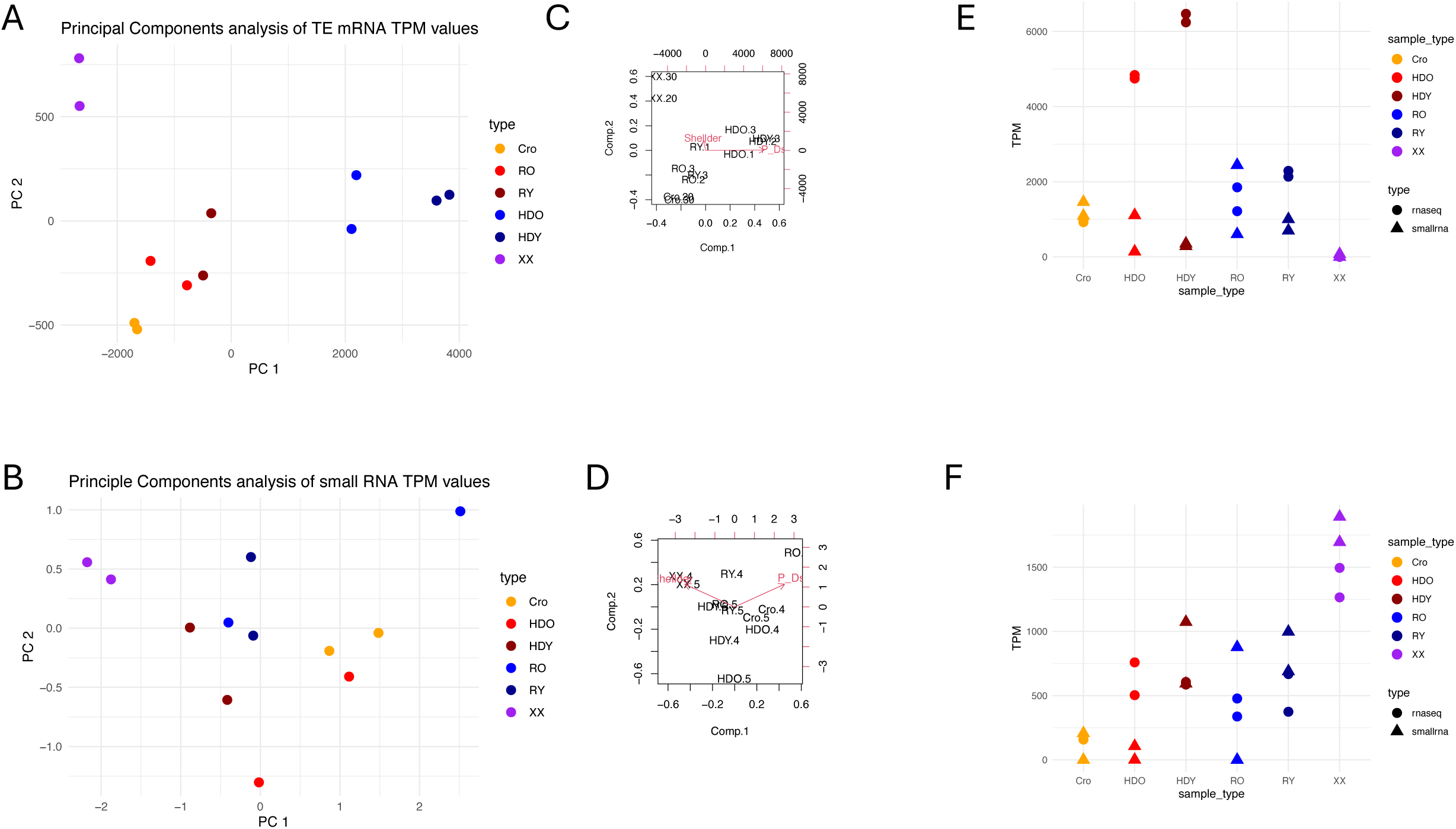
Principal components analyses of TPM values from SalmonTE on TE expression levels (A), piRNA expression levels (B) and bi-plots showing the major contributors to differences on PC1 (the P-element; P_Dsim) and PC2 (Shellder) for both TE mRNA (C) and piRNAs (D). Expression levels of P-element (E) and Shellder (F) in the different samples are shown.

**Supplementary Table 1.** Fly crosses and pooling strategy for transposition assays, used to measure transposition in the germline of dysgenic (HD) and reciprocal control (R) F1 males and females. Libraries targeting specific TEs (*P-*element and *roo*) were prepared from F2 flies produced as detailed below. Library indicates the type of individual targeted for transposition measurement. Pooled F2s are the sequenced flies. Crosses in bold are M-female x P-male crosses that may induce dysgenesis.

| Library | Initial cross<br>(28.5°C) | Second cross<br>(25°C) | Pools |  |  |
| --- | --- | --- | --- | --- | --- |
|  |  |  | sex | pools | flies/<br>pool |
| Cro18 | - | - | F | 1 | 8 |
|  | - | - | M | 1 | 8 |
| M252 | - | - | F | 1 | 8 |
| XX | - | - | Mixed | 1 | 8 |
| HD-♀ | **M252 ♀ x Cro18 ♂** | F1♀ x M252♂ | F | 59 | 472 |
| R-♀ | Cro18 ♀ x M252 ♂ | F1♀ x M252♂ | F | 61 | 488 |
| HD-♂ | **XX ♀ x Cro18 ♂** | **F1♂ x XX♀** | F | 29 | 232 |
|  |  |  | M | 41 | 328 |
|  |  |  | Mixed | 1 | 6 |
|  |  |  |  | *total* | *566* |
| R-♂ | Cro18 ♀ x XX ♂ | **F1♂ x XX♀** | F | 19 | 152 |
|  |  |  | M | 51 | 410 |
|  |  |  | Mixed | 1 | 8 |
|  |  |  |  | *total* | *570* |

**Supplementary Table 2.**
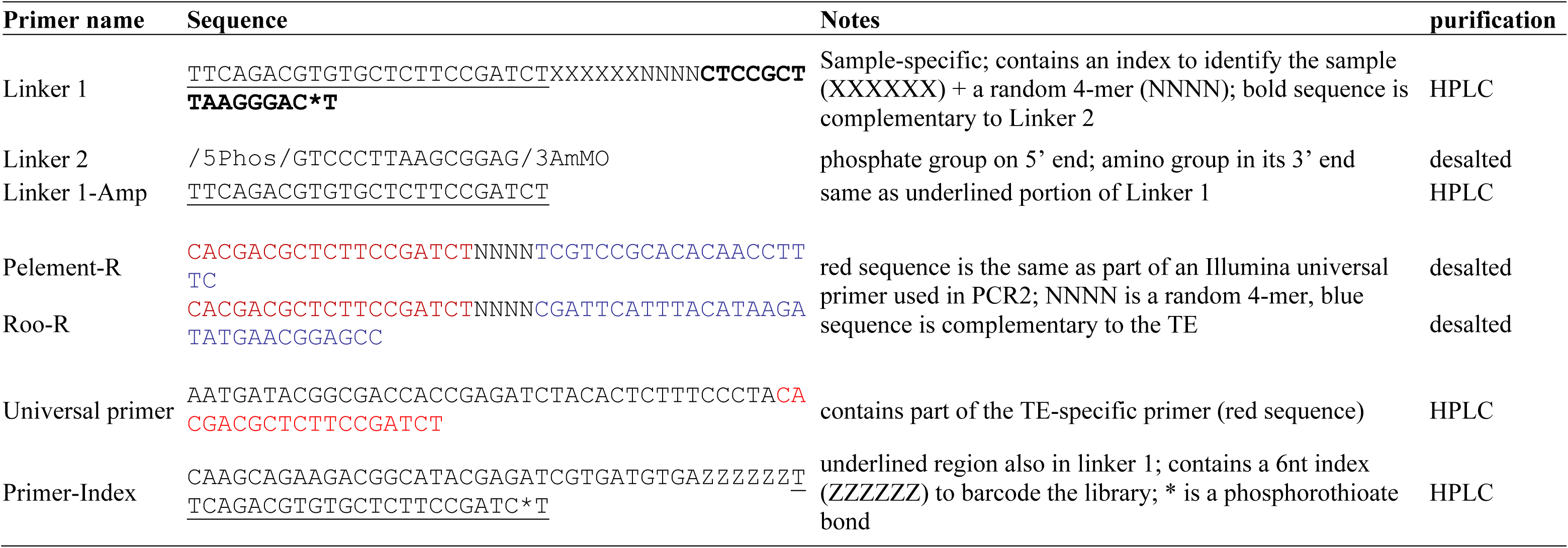
Oligos used in library preparation to identify TE insertion locations.

**Supplementary Table 3.** Characteristics of the RNAseq samples from testes used in this study. Mapping was done as described in the main text, and mapping rates are from Salmon or SalmonTE as appropriate. Sample names indicate whether the sample is from a parental strain (P-type Cro18 or M-type attached-X), or from an F1 and in which direction (HD = hybrid dysgenic, with an attached-X maternal parent, R = reciprocal, with a Cro18 maternal parent). Numbers are used to separate replicates.

| **Cross direction** | **Age** | **Sample** | **Library size**<br>**(n read pairs)** | **Mapping rate:**<br>**transcriptome** | **Mapping**<br>**rate: TEs** |
| --- | --- | --- | --- | --- | --- |
| **Parent** | Young | 20-Cro18 | 78,877,802 | **71.35%** | 0.5268 % |
| **Parent** | Young | 30-Cro18 | 76,828,078 | 72.07 | 0.5493 |
| **Parent** | Young | 20-XX | 69,919,986 | 87.01 | 0.6673 |
| **Parent** | Young | 30-XX | 85,535,342 | 86.77 | 0.6835 |
| **Dysgenic** | Young | 5-HDY | 85,556,986 | 83.46 | 0.5491 |
| **Dysgenic** | Young | 6-HDY | 87,834,072 | 85.80 | 0.4774 |
| **Dysgenic** | Old | 7-HDO | 69,779,714 | 82.87 | 0.8228 |
| **Dysgenic** | Old | 8-HDO | 97,114,280 | 82.40 | 1.0258 |
| **Reciprocal** | Young | 9-RY | 78,278,758 | 60.11 | 0.2772 |
| **Reciprocal** | Young | 10-RY | 60,430,212 | 65.74 | 0.3541 |
| **Reciprocal** | Old | 11-RO | 93,116,912 | 71.45 | 0.4289 |
| **Reciprocal** | Old | 12-RO | 73,876,468 | 75.56 | 0.7183 |

**Supplementary Table 4.** Estimates of the number of transcripts corresponding to splice variants of the P-element, resulting in transcripts encoding transposase (IVS3 spliced; coordinates in Genbank accession X06779: 1947-2138) or one of two alternatively spliced transcripts encoding a truncated protein. Estimates of splicing of this intron are STAR as described in the main text. IVS3 stands for intervening sequence 3, or the third intron of the P-element transcript. In IV3 spliced transcripts, this intron is removed fully. IV3.1 and IV3.2 are subsequences of this intron that spliced from alternate donor (GT-) and acceptor sites (-AG), resulting in the retention of a small exon (at 2015-2049), resulting in a transcript encodingg a truncated transposase; the ‘spliced non-functional’ column contains the estimated number of transcripts containing this exon from the average of IV3.1 and IVS3.2 splice estimates. The columns Exon2-IVS3 and IVS3-Exon3 junctions are obtained from reads covering the intron-exon junctions from samtools depth, and are averaged to estimate the number of unspliced transcripts. The sum of spliced non-functional and unspliced transcripts is summed for the total number of non-functional transcripts. Numbers in bold are those used for statistical tests of functional *vs* nonfunctional transcripts for each sample type.

| Sample | Coding transcript | Transcripts encoding non-functional transposase |  |  |  |  |  |  |
| --- | --- | --- | --- | --- | --- | --- | --- | --- |
|  | IVS3 spliced | IVS3.1 | IVS3.2 | spliced non-functional | Exon2-IVS3 junction | IVS3-Exon3 junction | fully unspliced | All non-functional |
| P-type (Cro18) | **0** | 3 | 1 | 2 | 28 | 46 | 37 | **39** |
| P-type (Cro18) | **0** | 2 | 1 | 2 | 32 | 25 | 28.5 | **30** |
| Hybrid Dysgenic Old | **8** | 83 | 76 | 80 | 127 | 146 | 136.5 | **216** |
| Hybrid Dysgenic Old | **5** | 55 | 50 | 53 | 162 | 169 | 165.5 | **218** |
| Hybrid Dysgenic Young | **12** | 62 | 64 | 63 | 114 | 177 | 145.5 | **209** |
| Hybrid Dysgenic Young | **7** | 52 | 49 | 51 | 130 | 142 | 136 | **187** |
| Reciprocal Old | **0** | 20 | 18 | 19 | 31 | 43 | 37 | **56** |
| Reciprocal Old | **0** | 8 | 7 | 8 | 37 | 35 | 36 | **44** |
| Reciprocal Young | **0** | 3 | 4 | 4 | 29 | 30 | 29.5 | **33** |
| Reciprocal Young | **0** | 11 | 6 | 9 | 26 | 23 | 24.5 | **33** |
| M-type (attached X) | **0** | 0 | 0 | 0 | 0 | 0 | 0 | **0** |
| M-type (attached X) | **0** | 0 | 0 | 0 | 0 | 0 | 0 | **0** |

**Supplementary Table 5.** Significant GO terms for MF and CC in the 112 upregulated DEGS for dysgenic vs. reciprocal flies that were common to the testes of young and old flies.

|  | **GO.ID** | **Term** | **Annotated** | **Significant** | **Expected** | **classicFisher** |
| --- | --- | --- | --- | --- | --- | --- |
| Molecular Function | GO:0050839 | cell adhesion molecule binding | 17 | 3 | 0.11 | 0.00016 |
|  | GO:1901363 | heterocyclic compound binding | 786 | 11 | 5.05 | 0.0079 |
| Cellular Component | GO:0045178 | basal part of cell | 35 | 3 | 0.2 | 0.00099 |
|  | GO:0098858 | actin-based cell projection | 12 | 2 | 0.07 | 0.00203 |
|  | GO:0098590 | plasma membrane region | 124 | 4 | 0.71 | 0.00516 |
|  | GO:0009925 | basal plasma membrane | 25 | 2 | 0.14 | 0.00881 |

**Supplementary Table 6.** Significant GO terms for BF, MF and CC in the 104 downregulated DEGS for dysgenic vs. reciprocal flies that were common to the testes of young and old flies.

|  | **GO.ID** | **Term** | **Annotated** | **Significant** | **Expected** | **classicFisher** |
| --- | --- | --- | --- | --- | --- | --- |
| Biological Function | GO:0000413 | protein peptidyl-prolyl isomerization | 16 | 2 | 0.04 | 0.00085 |
|  | GO:0018208 | peptidyl-proline modification | 16 | 2 | 0.04 | 0.00085 |
| Molecular Function | GO:0003755 | peptidyl-prolyl cis-trans isomerase acti... | 27 | 2 | 0.06 | 0.0016 |
|  | GO:0016859 | cis-trans isomerase activity | 29 | 2 | 0.07 | 0.0019 |
| Cellular Component | GO:0016020 | membrane | 2543 | 12 | 7.06 | 0.0098 |

**Supplementary Table 7.** Characteristics of the small RNAseq samples from testes used in this study. Filtering and mapping were done as described in the main text. Sample names indicate whether the sample is from a parental strain (P-type Cro18 or M-type attached-X), or from an F1 and in which direction (HD = hybrid dysgenic, with an attached-X maternal parent, R = reciprocal, with a Cro18 maternal parent). Numbers are used to separate replicates.

| **Cross direction** | **Age** | **Sample** | **Library size pre-filtering** | **Library size post-filtering** | **SalmonTE mapping rate** |
| --- | --- | --- | --- | --- | --- |
| **Parent** | Young | 13-Cro-4 | 20,336,238 | 7,295,922 | 0.1544 % |
| **Parent** | Young | 14-Cro-5 | 22,489,472 | 8,020,712 | 0.1707 |
| **Parent** | Young | 15-XX-4 | 18,617,008 | 6,550,881 | 0.1563 |
| **Parent** | Young | 16-XX-5 | 61,032,013 | 22,239,761 | 0.1305 |
| **Dysgenic** | Young | 17-HDY | 9,727,359 | 3,227,352 | 0.1082 |
| **Dysgenic** | Young | 18-HDY | 10,252,686 | 3,876,066 | 0.1700 |
| **Dysgenic** | Old | 19-HDO | 4,086,151 | 1,723,350 | 0.1980 |
| **Dysgenic** | Old | 20-HDO | 87,242,240 | 39,386,956 | 0.2314 |
| **Reciprocal** | Young | 21-RY | 31,472,367 | 12,916,110 | 0.1496 |
| **Reciprocal** | Young | 22-RY | 65,356,370 | 25,511,053 | 0.1161 |
| **Reciprocal** | Old | 23-RO | 1,122,033 | 396,804 | 0.1285 |
| **Reciprocal** | Old | 24-RO | 16,417,402 | 5,420,735 | 0.1409 |

